# Kinetic control of mammalian transcription elongation

**DOI:** 10.1101/2025.02.24.639907

**Authors:** Yukun Wang, Xizi Chen, Maximilian Kuemmecke, John W. Watters, Joel E. Cohen, Yanhui Xu, Shixin Liu

## Abstract

Transcription elongation by RNA polymerase II is an integral step in eukaryotic gene expression. The speed of Pol II is controlled by a multitude of elongation factors, but the regulatory mechanisms remain incompletely understood, especially for higher eukaryotes. In this work, we developed a single-molecule platform to visualize the dynamics of individual mammalian transcription elongation complexes (ECs) reconstituted from purified proteins. This platform enabled us to follow the elongation and pausing behavior of EC in real time and dissect the role of each elongation factor in the kinetic control of Pol II. We found that the mammalian EC harbors multiple gears dictated by its associated factors and phosphorylation status. Moreover, the elongation factors are not functionally redundant but act hierarchically and synergistically to achieve optimal elongation activity. Such exquisite kinetic regulation may underline the major speed-changing events during the transcription cycle and enable cells to adapt to a changing environment.

## Introduction

Transcription elongation by RNA polymerase II (Pol II) is being increasingly recognized to be a focal point of regulation in eukaryotic gene expression,^1,2^ and its dysregulation has been linked to disease and aging.^3,4^ In metazoans, after Pol II assembles the initiation complex at the promoter and transcribes 50-100 nucleotides (nt) of RNA, it enters a paused state known as promoter-proximal pausing.^5^ Several elongation factors, including P-TEFb, DSIF and PAF1C, control the release of Pol II into productive elongation.^6^ At this stage, Pol II speeds up to several kilobases (kb) per minute, reads through exons and introns, and is kinetically coupled with other co-transcriptional processes such as messenger RNA (mRNA) splicing.^7,8^ At the 3’ end of genes, Pol II slows down to facilitate co-transcriptional mRNA cleavage and polyadenylation, events critical for the proper termination of transcription.^9^

The activity of the mammalian elongation complex (EC) is regulated by myriad elongation factors and complexes, which decorate a major fraction of the Pol II surface as revealed by structural studies (Figure 1A).^10,11^ Much of the knowledge regarding the functions of these regulatory factors came from in vivo studies, particularly those using genetic or chemical perturbations.^12–14^ In vitro biochemical studies offer a complementary approach to determine whether a factor is stimulatory or repressive to EC activity.^15–17^ However, the readouts of these assays are either stable genomic positions of EC components or endpoint RNA products from many transcription events. It remains challenging to distinguish direct versus indirect effects on the elongation activity by specific factors from often subtle phenotypes. Filling this knowledge gap requires assays capable of following the elongation kinetics in real time and directly relating factors binding to EC activity.

**Figure 1.**
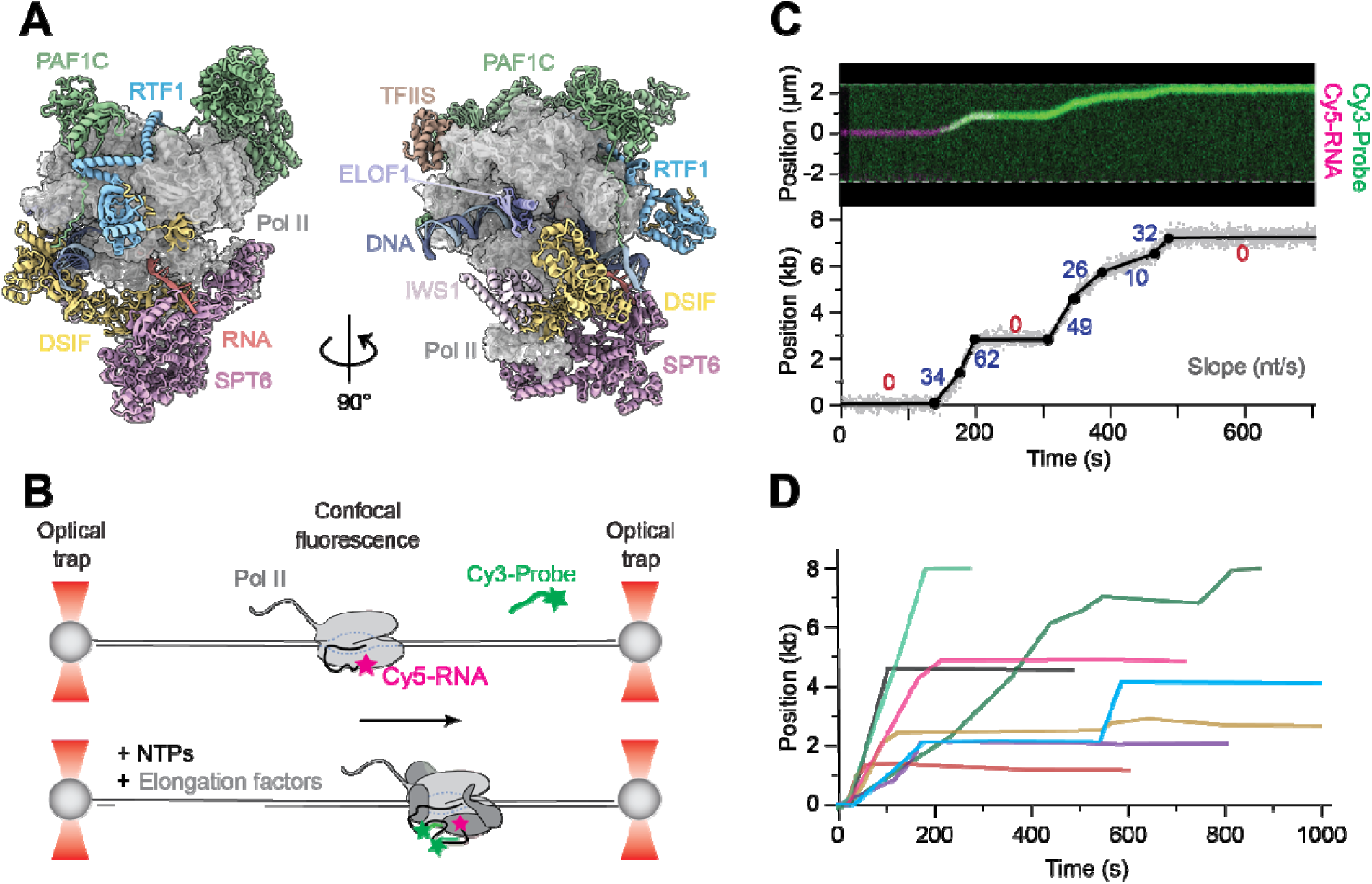
Real-time observation of mammalian transcription elongation. (A) Composite structural model of the human transcription elongation complex (EC) with a set of elongation factors, as supplied in our system, decorating the Pol II surface. The model incorporates published structures for the mammalian EC (PDB: 6TED), TFIIS (PDB: 8A40), ELOF1 (PDB: 8B3F), and a fitted AlphaFold 2 prediction of IWS1 based on the position of Spn1 in the yeast EC (PDB: 7XN7). (B) Schematic of the single-molecule experimental setup. The starting location of the EC is marked by the Cy5-labeled RNA, while Cy3-labeled DNA probes that hybridize to the nascent RNA indicate active elongation. (C) An example of transcription elongation by a single mammalian EC observed in real time. (Top) Kymograph showing the EC position on the DNA template, indicated by Cy5-RNA (magenta) and Cy3-probe (green) signals, as a function of time. (Bottom) Elongation trajectory extracted from the kymograph. Raw data (gray dots) were fitted to discrete linear segment (black line). Changepoints are marked as filled circles. The slopes for each segment are indicated, differentiating active elongation (blue) from pausing events (red). (D) 8 additional examples of fitted elongation trajectories by individual mammalian ECs aligned by their starting positions.

Single-molecule techniques circumvent the limitations associated with ensemble averaging, thus well suited for studying dynamic biomolecular processes.^18,19^ Indeed, single-molecule studies have provided a wealth of insights into the mechanism and regulation of transcription elongation in model systems such as *Escherichia coli* and *Saccharomyces cerevisiae*.^20,21^ But a similar level of mechanistic understanding has not been achieved for higher eukaryotic systems, largely due to the increased difficulty in reconstituting active ECs at the single-molecule level. The only previous study to our knowledge reported a low basal activity of the mammalian Pol II compared to the yeast counterpart^22^, indicating that certain elongation factors were missing from the study.

In this work, we reconstituted the 26-subunit mammalian EC with a total molecular weight of 1.6 megadalton using purified proteins at the single-molecule level. This allowed us to measure the real-time elongation and pausing kinetics of mammalian transcription and dissect the interplay between elongation factors in the kinetic control of Pol II. Our results elucidated the hierarchical roles of several key elongation factors including P-TEFb, DSIF, PAF1C, SPT6, and RTF1, and provide a quantitative framework for understanding the intricate regulation of mammalian transcription elongation.

## Results

### Single-molecule visualization of mammalian transcription elongation

We purified the mammalian Pol II (Figure S1A) and assembled EC using a synthetic nucleic acid scaffold containing a fluorescently labeled RNA primer and template/non-template DNA which was pre-ligated to two 8-kb-long biotinylated DNA handles on either side (Figure S2). The 16-kb DNA with an EC positioned in the middle was tethered between a pair of streptavidin-coated beads and visualized on a C-Trap instrument that combines dual-trap optical tweezers and single-molecule fluorescence microscopy (Figure 1B).^23^ For enhanced fluorescence detection, the downstream DNA contained seven repeat sequences that, when transcribed into nascent RNA, could each be hybridized to a fluorescently labeled DNA probe. When supplied with eight purified human elongation factors/complexes, i.e. P-TEFb, DSIF, PAF1C, RTF1, SPT6, IWS1, TFIIS, and ELOF1 (Figure S1A) at saturating concentrations, more than half of the ECs exhibited active elongation, indicated by the movement of the Cy5-RNA fluorescence signal and the appearance/movement of the Cy3-probe fluorescence signal (Figures 1C and S3).

We found mammalian transcription elongation to be a highly dynamic and heterogeneous process, frequently transitioning between different kinetic states. To quantitatively characterize the kinetic behavior of the EC, we developed a Bayesian framework to divide individual elongation trajectories into discrete linear segments. We defined the segments with a positive slope of ≥1 nt/s as active elongation, and those with a slope of <1 nt/s or negative values as pausing events. The observed transient pauses may result from Pol II arrest or backtracking, which can be rescued by the elongation factor TFIIS that induces RNA cleavage.^24,25^ Indeed, when we omitted TFIIS from the single-molecule assay, we saw a significant increase in the pause duration (Figure S4A). Moreover, we found that most EC trajectories ended with a long-lasting stall that featured extensive backtracking (Figures 1D and S4B). Unlike the transient pauses, these stalling events cannot be resolved by TFIIS (Figure S4C), potentially corresponding to the “persistent backtracking” phenomenon found in mammalian cells.^26^ It remains to be determined which additional factor(s) are required to resolve the long-lasting stall for Pol II to transcribe long genes in vivo.

### Elongation factors exert differential effects on EC activity

Next, we focused on the active elongation segments, whose slopes exhibit a wide distribution that spans two orders of magnitude (Figure 2A). When all factors were included, the average pause-free speed, weighted by the transcribed length of each active segment, was 40 ± 1 nt/s (Figure 2B). We then recorded single-molecule trajectories under conditions where individual elongation factors were omitted, which allowed us to determine their respective contribution to the elongation kinetics (Figures 2A and S5). The single-omission experiments yielded a spectrum of phenotypes. Some conditions (such as ΔP-TEFb and ΔPAF1C) exhibited a drastic reduction in speed, while others (such as ΔIWS1) showed only a mild effect. Interestingly, when plotting the average speed against the average distance traveled before stalling for each condition, we found that most of the conditions followed a linear relationship (Figure 2B). This indicates that the total time an EC spends on productive elongation (distance over speed) remained largely constant across different conditions. However, there were two notable exceptions: ΔDSIF resulted in significantly reduced speed with a slight decrease in distance, whereas ΔSPT6 drastically shortened the distance but only moderately affected the pause-free speed. In the following, we will investigate the role of those factors that displayed a strong phenotype in the control of elongation kinetics, namely P-TEFb, DSIF, PAF1C, SPT6, and RTF1.

**Figure 2.**
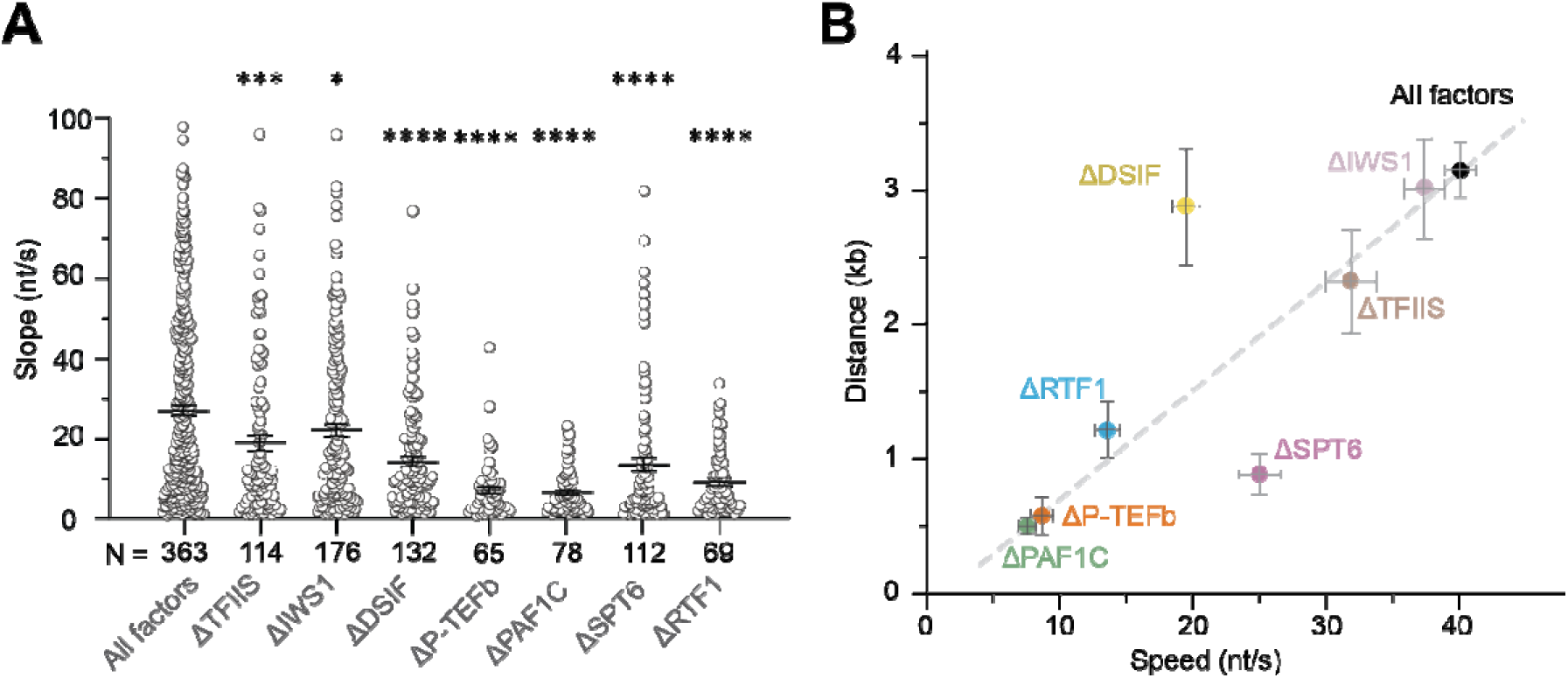
Differential effects of elongation factors on EC kinetics. (A) Dot plot showing the slope of active elongation segments from EC trajectories under conditions where individual elongation factors were omitted. Bars represent mean ± SEM. \**P* < 0.05; \*\*\**P* < 0.001; \*\*\*\**P* < 0.0001. *P* values were from unpaired t tests with Welch’s correction against the All factors condition. (B) The average distance traveled before stalling by individual mammalian ECs at a given condition (normalized by the active EC fraction) is plotted against the average pause-free elongation speed weighted by the length of the active segments. Error bars represent SEM. The dashed line represents the linear regression of the speed vs. distance values for all conditions.

### P-TEFb activates elongation by phosphorylating Pol II and DSIF

P-TEFb is a positive elongation factor that contains a cyclin-dependent kinase CDK9, which phosphorylates the C-terminal domain (CTD) of Pol II’s RPB1 subunit, as well as DSIF, PAF1C and SPT6.^10,16,27,28^ Among these phosphorylated substrates, which one(s) directly impact the elongation speed remains unclear. In our single-molecule assay, omitting P-TEFb reduced the elongation speed and distance by ∼5-fold (compare ΔP-TEFb to All factors in Figure 2B). To separate the effects of multiple CDK9-mediated phosphorylation products on the elongation kinetics, we specifically prevented the phosphorylation of either Pol II or DSIF (STAR Methods) and found that both Pol II^P-^ and DSIF^P-^ conditions exhibited a severe elongation defect (Figures 3A-3D and S6). We then performed experiments in which only Pol II and DSIF—but not any other factor—were allowed to be phosphorylated and found that this condition (Pol II^P+^&DSIF^P+^) phenocopied the condition where all factors were allowed to be phosphorylated by P-TEFb (Figures 3A, 3B, 3E, and S6). Together, these results suggest that the phosphorylation of Pol II and DSIF is both necessary and sufficient for optimal EC activity.

**Figure 3.**
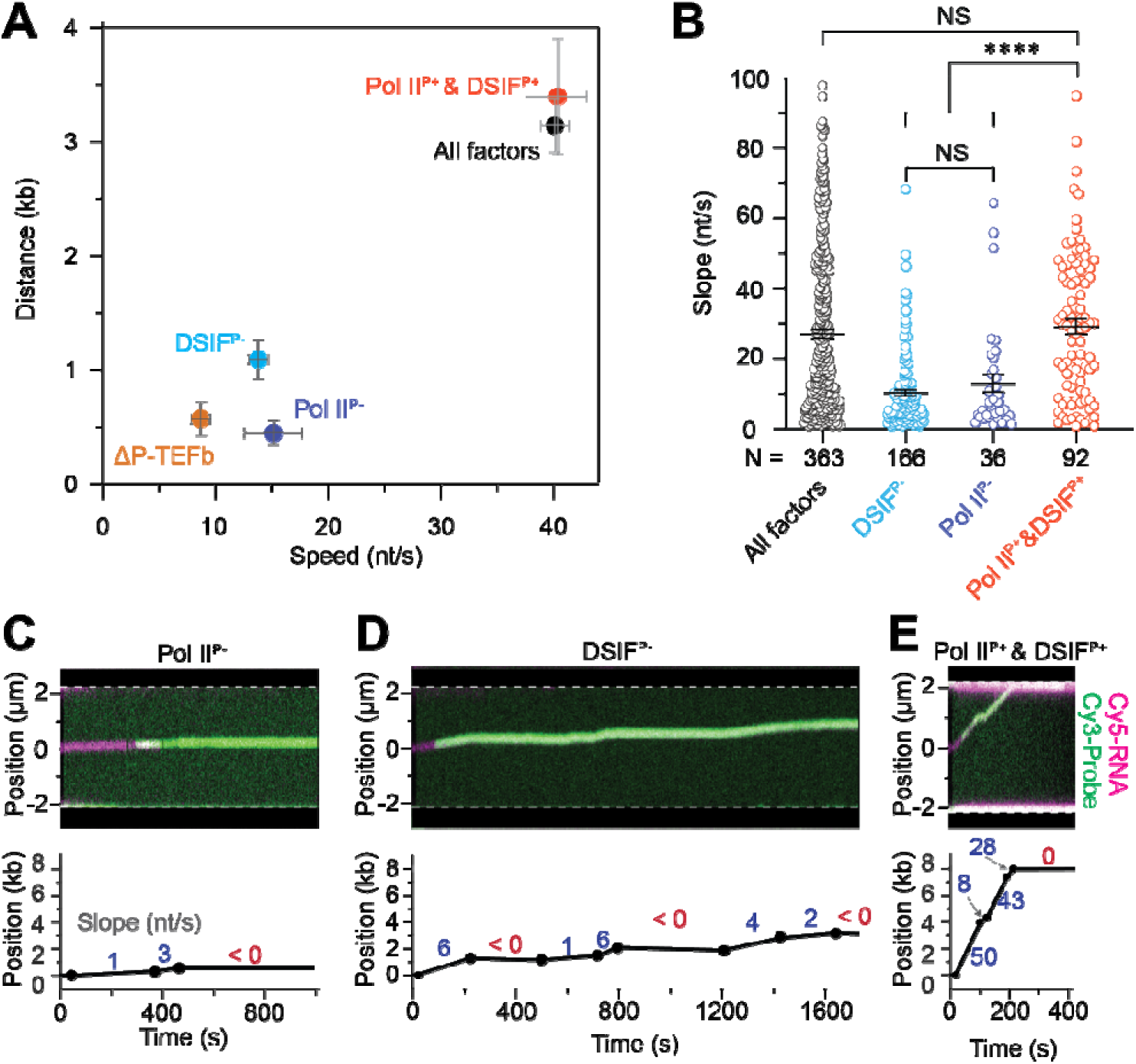
Effects of P-TEFb-mediated phosphorylation on EC activity. (A) 2D plot showing the average elongation speed and distance for different phosphorylation conditions. All factors: the full set of factors present, all of which were allowed to be phosphorylated by P-TEFb; ΔP-TEFb: P-TEFb omitted, hence no phosphorylation on any protein; DSIF^P-^: all factors present, but phosphorylation of DSIF was specifically inhibited; Pol II^P-^: all factors present, but Pol II was not phosphorylated; Pol II^P+^&DSIF^P+^: all factors present, but only Pol II and DSIF were allowed to be phosphorylated. Error bars represent SEM. (B) Dot plot showing the slope of active elongation segments from EC trajectories under All factors, DSIF^P-^, Pol II^P-^, and Pol II^P+^&DSIF^P+^ conditions. Bars represent mean ± SEM. NS, not significant; \*\*\*\**P* < 0.0001. *P* values were from unpaired t tests with Welch’s correction. (C) An example EC trajectory for the Pol II^P-^ condition. (Top) Kymograph showing the overlay of Cy5-labeled RNA (magenta) and Cy3-labeled complementary DNA probes (green) which were used to track EC progression. (Bottom) Fitted and segmented trajectory with the slope of each segment indicated. (D) An example EC trajectory for the DSIF^P-^ condition. Details are the same as (C). (E) An example EC trajectory for the Pol II^P+^&DSIF^P+^ condition. Details are the same as (C).

### DSIF regulates both elongation and pausing kinetics of EC

DSIF exerts pleiotropic effects on transcription in vivo.^13,29–31^ In vitro biochemical assays also suggest that DSIF can both negatively and positively regulate elongation.^32,33^ To gain insight into the behavior of DSIF during elongation, we fluorescently labeled its SPT4 subunit and employed two-color imaging to simultaneously monitor DSIF binding and EC translocation. DSIF and EC were frequently observed to colocalize and comigrate on the DNA template (Figures 4A and S7A), suggesting stable association of DSIF with Pol II during elongation. We also observed non-EC-bound DSIF on DNA, often exhibiting a diffusive behavior (Figure 4A). The biological significance of such Pol II-independent DNA binding by DSIF awaits further investigation. We then used the CDK9 inhibitor flavopiridol to inhibit DSIF phosphorylation by P-TEFb and imaged its interaction with EC (Figures 4B and S7B). We observed that unphosphorylated DSIF binds EC much more transiently compared to the phosphorylated version (47 ± 4 s for DSIF^P-^ vs. 182 ± 14 s for DSIF^P+^) (Figure 4C), revealing that stable association of DSIF with EC requires phosphorylated DSIF. Notably, unphosphorylated DSIF also loses the ability to maintain fast elongation (Figures 4B and S7B).

**Figure 4.**
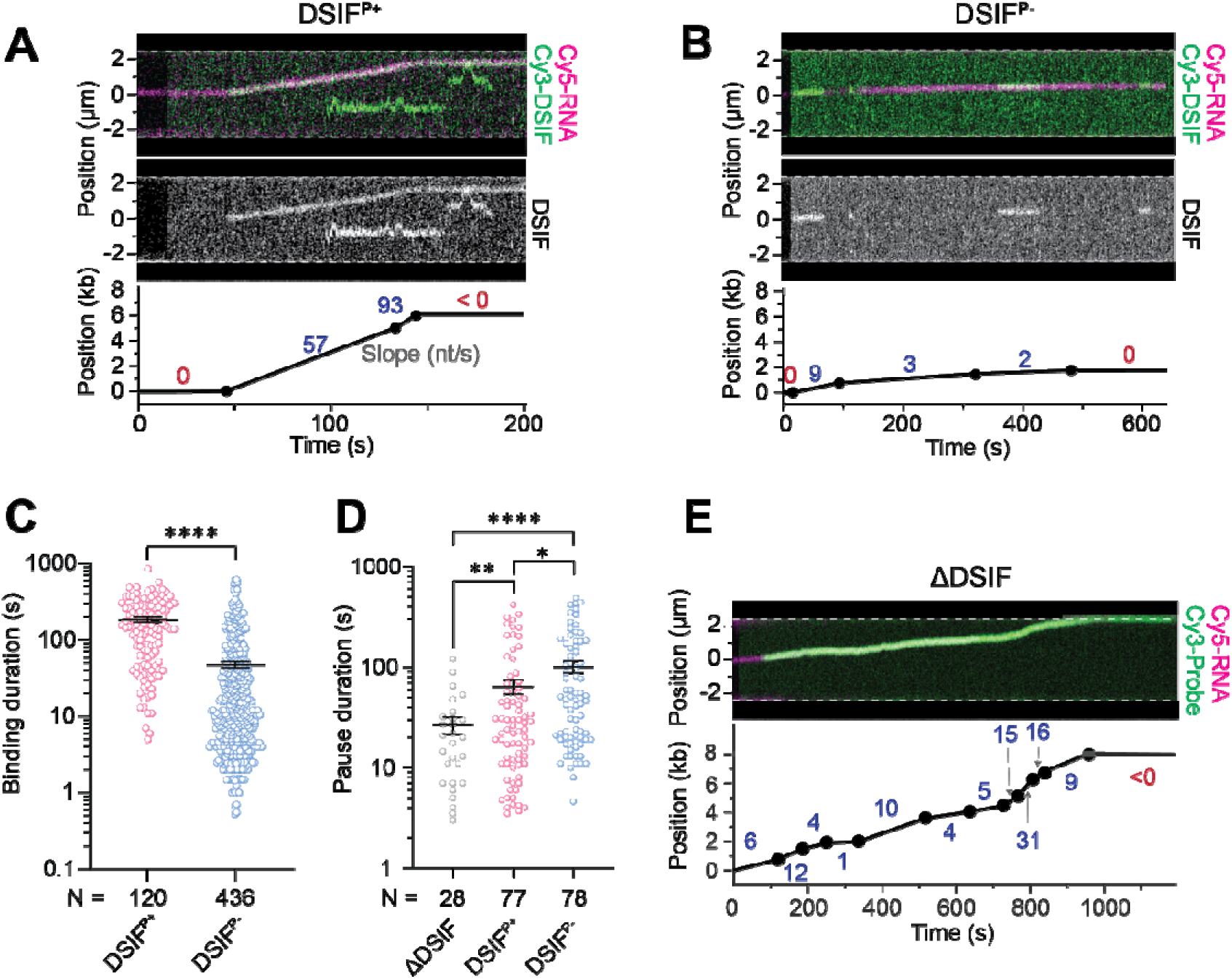
DSIF plays a dual role in EC regulation. (A) An example EC trajectory with Cy3-labeled phosphorylated DSIF (DSIF^P+^) and Cy5-labeled RNA. (Top) Kymograph showing the overlay of DSIF (green) and RNA (magenta) signals. (Middle) The same kymograph but with only DSIF signals shown to more clearly visualize its binding to the EC and DNA. (Bottom) Fitted and segmented elongation trajectory with the slope of each segment indicated (active elongation in blue and pausing/stalling in red). (B) An example EC trajectory with Cy3-labeled unphosphorylated DSIF (DSIF^P-^). Details are the same as (A). (C) Dot plot showing the lifetime of DSIF^P+^ (red) and DSIF^P-^ (blue) binding events on EC. \*\*\*\**P* < 0.0001. (D) Dot plot showing the duration of pauses within the elongation trajectories under th condition where DSIF was omitted (ΔDSIF) or included as either a phosphorylated (DSIF^P+^) or an unphosphorylated (DSIF^P-^) form. \**P* < 0.05; \*\**P* < 0.01; \*\*\*\**P* < 0.0001. *P* values in (C) and (D) were from unpaired t tests with Welch’s correction. Bars represent mean ± SEM. (E) An example EC trajectory for the ΔDSIF condition. (Top) Kymograph showing the overlay of Cy5-labeled RNA (magenta) and Cy3-labeled complementary DNA probes (green) which were used to track EC progression. (Bottom) Fitted and segmented trajectory with the slope of each segment indicated.

We found that the omission of DSIF reduced the EC speed by ∼2-fold but minimally affected the distance it can travel (compare ΔDSIF to All factors in Figure 2B). The deviation from the linear speed-distance relationship indicates that the fraction of time an EC spends on productive elongation is affected by DSIF. We thus analyzed the pausing kinetics and indeed found that EC spent significantly less time in a paused state when DSIF was omitted (Figures 4D, 4E, and S7C). This finding reveals that besides its positive effect on the elongation speed, DSIF also exerts a pause-stabilizing effect on the EC, possibly related to its DNA and RNA clamping activity.^34,35^ As such, the reduced pausing under the ΔDSIF condition allows Pol II to travel for similar distances before stalling, albeit at a reduced speed. Moreover, when used at saturating concentrations, DSIF^P+^ and DSIF^P-^ both caused longer pauses compared to the ΔDSIF condition (Figure 4D), indicating that DSIF promotes pausing regardless of its phosphorylation status.

### PAF1C binding directly accelerates EC

PAF1C has been shown to promote active elongation in vitro^10,15^ and in vivo.^14,36–38^ We found that omitting PAF1C caused severely diminished elongation speed and distance in our single-molecule assay (Figures 2B and S8A). To directly visualize the behavior of PAF1C, we attached a Cy3 fluorophore to its CTR9 subunit and monitored its interaction with EC during elongation (Figures 5A and S8B). Strikingly, we observed that in the majority (71%) of active trajectories, the onset of PAF1C signal coincided with a sudden acceleration of the EC (i.e. Δ*t* = *t*_move_ - *t*_bind_ = 0), indicating an immediate stimulatory effect of PAF1C association on elongation. In comparison, DSIF association and EC speed change co-occurred in only 29% of the trajectories. We also observed that the PAF1C fluorescence signal continued to increase in a stepwise manner as elongation progressed (Figure S8C). We speculated that additional copies of PAF1C may be recruited to the growing RNA chain as shown previously in the yeast system.^39^ Indeed, RNase A treatment led to a constant level of PAF1C signals throughout the trajectory, suggesting the association of one single copy of PAF1C with the EC without RNA (Figure S8D). This treatment allowed us to measure the lifetime of PAF1C-Pol II interaction to be 80 ± 10 s (mean ± SEM, *N* = 128). Our in vitro data thus show that the residence time of PAF1C on the EC is shorter than that of DSIF, consistent with recent live-cell imaging results.^40^

**Figure 5.**
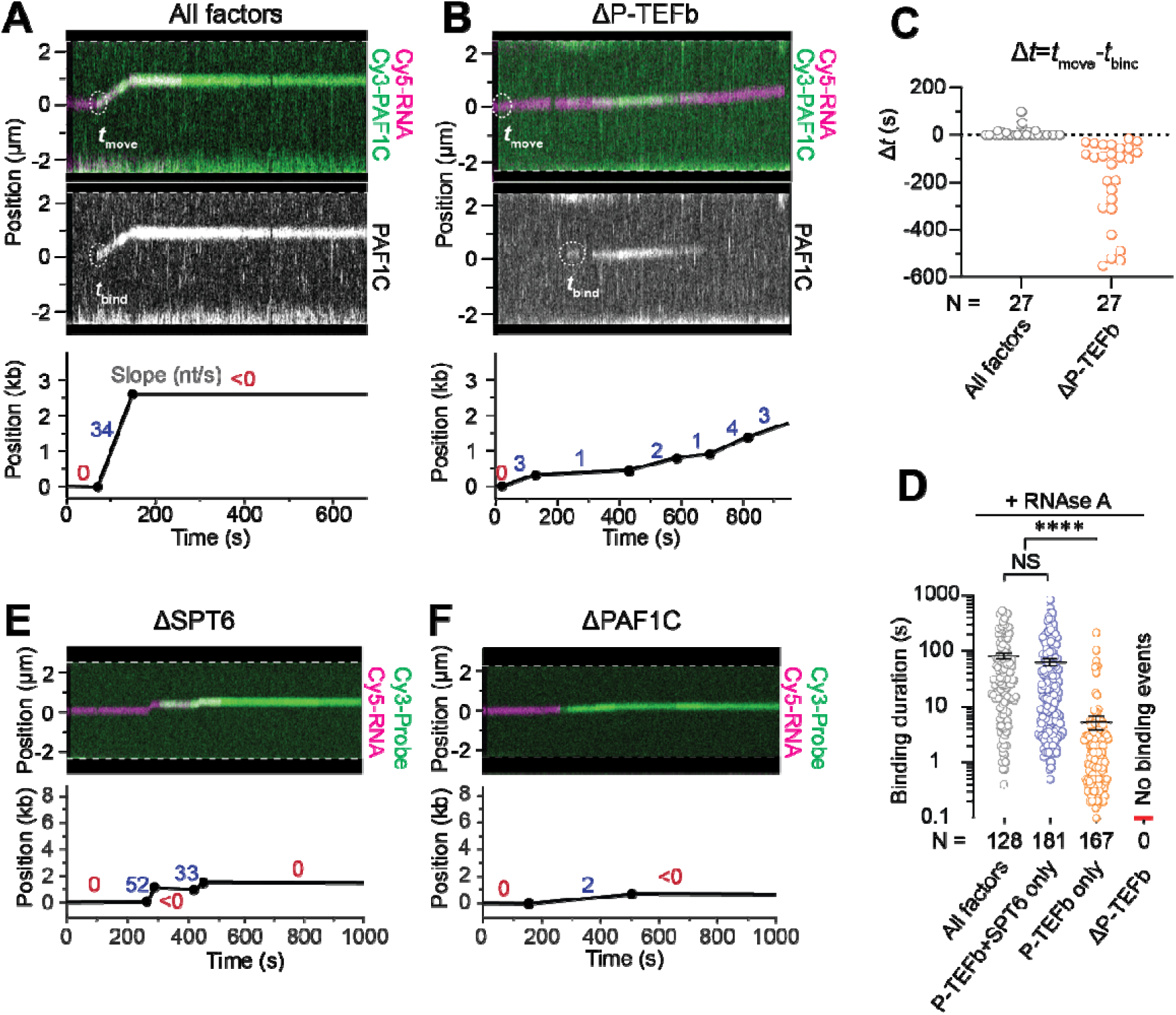
PAF1C binding directly activates EC and is stabilized by SPT6. (A) An example EC trajectory with Cy3-labeled PAF1C and Cy5-labeled RNA when the full set of elongation factors were present (All factors). (Top) Kymograph showing the overlay of PAF1C (green) and RNA (magenta) signals. *t*_move_ indicates the time point when the EC started translocation. (Middle) The same kymograph but with only PAF1C signals shown. *t*_bind_ indicates the time point when PAF1C binding to EC occurred. (Bottom) Fitted and segmented trajectory with the slope of each segment indicated (active elongation in blue and pausing/stalling in red). (B) An example EC trajectory with Cy3-PAF1C and Cy5-RNA when P-TEFb was omitted (ΔP-TEFb). Details are the same as (A). (C) Dot plot showing the Δ*t* value (Δ*t* = *t*_move_ - *t*_bind_) for individual EC trajectories from All factor (black) and ΔP-TEFb (orange) conditions. (D) Dot plot showing the lifetime of PAF1C binding events on EC under different conditions. RNase A was included in these experiments to eliminate RNA-mediated PAF1C recruitment. Bars represent mean ± SEM. NS, not significant; \*\*\*\**P* < 0.0001. *P* values were from unpaired t tests with Welch’s correction. (E) An example EC trajectory under the ΔSPT6 condition. (Top) Kymograph showing the overlay of Cy5-labeled RNA (magenta) and Cy3-labeled complementary DNA probes (green) which were used to track EC progression. (Bottom) Fitted and segmented trajectory with the slope of each segment indicated. (F) An example EC trajectory under the ΔPAF1C condition. Details are the same as (E).

Given that the ΔPAF1C and ΔP-TEFb conditions essentially phenocopied each other (Figure 2B), we asked whether P-TEFb is responsible for PAF1C recruitment to EC. To this end, we monitored PAF1C binding and EC translocation in the absence of P-TEFb (Figures 5B and S8E) and found that the appearance of the PAF1C signal was no longer coincident with EC movement (i.e. Δ*t* < 0; Figure 5C). Upon RNase A treatment, PAF1C binding was completely abolished (Figure 5D), suggesting that the PAF1C binding events under the ΔP-TEFb condition purely resulted from RNA-mediated recruitment. Collectively, these results reveal that PAF1C binding to Pol II—but not its binding to RNA—causally accelerates EC and this binding critically relies on P-TEFb, presumably via the interaction between phosphorylated CTD and the CDC73 subunit of PAF1C based on biochemical results from the yeast system.^41^

### SPT6 stabilizes PAF1C binding to EC

SPT6 has been shown to promote transcription elongation, which was attributed to its positive effect on the elongation rate and/or processivity.^12,14,42–44^ In our single-molecule assay, omitting SPT6 caused a moderately reduced elongation speed and a significantly shorter distance (ΔSPT6 in Figure 2B). Moreover, we observed that the EC trajectories under this condition frequently exhibited short elongation segments with relatively fast speeds interspersed with pauses (Figures 5E and S9A). This is different from the ΔPAF1C phenotype where the elongation segments per se were very slow (Figures 5F and S8A). Given previous in vivo depletion experiments suggesting that SPT6 helps recruit PAF1C to the EC,^12^ we examined the impact of SPT6 on PAF1C-Pol II interaction. We used fluorescently labeled PAF1C to visualize its binding in the presence of RNase A to eliminate RNA-mediated recruitment. When only P-TEFb and SPT6 were included, the residence time of PAF1C on the EC is statistically identical to the value when all factors were present (Figure 5D), suggesting that P-TEFb and SPT6 are sufficient for mediating stable PAF1C recruitment. However, further removing SPT6 caused a 10-fold reduction in the PAF1C residence time (Figure 5D). Overall, these results show that SPT6 itself does not greatly impact the pause-free speed; rather, it promotes elongation by stabilizing PAF1C-Pol II interaction.

Studies on the yeast system suggested an interaction between the SH2 domain of Spt6 and the Cdc73 subunit of Paf1C,^45,46^ but a structural link between PAF1C and SPT6 in the human EC has not been established.^10^ To explore how SPT6 helps stabilize PAF1C on the EC, we utilized AlphaFold 3 (ref. ^47^) to predict SPT6-PAF1C-Pol II interactions (Figure S9B). Our model suggests that the phosphorylated CTD of Pol II and SPT6 may simultaneously engage with PAF1C without steric hindrance (Figure S9C), corroborating our experimental results.

### RTF1’s role in EC stimulation differentially relies on PAF1C and DSIF

RTF1 is an essential elongation factor^11,14,17^ that makes extensive contacts with PAF1C and DSIF in the active EC structure^11^ (Figure 1A). Omitting RTF1 from our single-molecule experiments caused a significantly lower EC activity (Figures 6A and S10A), even though the phenotype was less severe compared to ΔPAF1C (Figure 2B). To further understand RTF1’s stimulatory function, we asked whether its effect on EC relies on the presence of PAF1C or DSIF, for which no consensus has been reached yet.^11,17^ To this end, we performed combinatorial omission experiments. First, we omitted both RTF1 and PAF1C and found that this condition (ΔRTF1&ΔPAF1C) yielded a phenotype more severe than ΔRTF1 but similar to ΔPAF1C (Figures 6A-6D, and S10B). The elongation speed for ΔRTF1&ΔPAF1C was significantly lower than that for ΔRTF1 but indistinguishable from that for ΔPAF1C (Figure 6B). Next, we omitted both DSIF and RTF1 and found that this condition (ΔRTF1&ΔDSIF) caused lower elongation speed and shorter transcription distance compared to either ΔRTF1 or ΔDSIF condition (Figures 6A, 6E, and S10C). These results suggest that PAF1C’s binding to the EC is a prerequisite for RTF1’s elongation-stimulating effect, and that RTF1 and DSIF can each exert an effect on the EC without the other’s presence.

**Figure 6.**
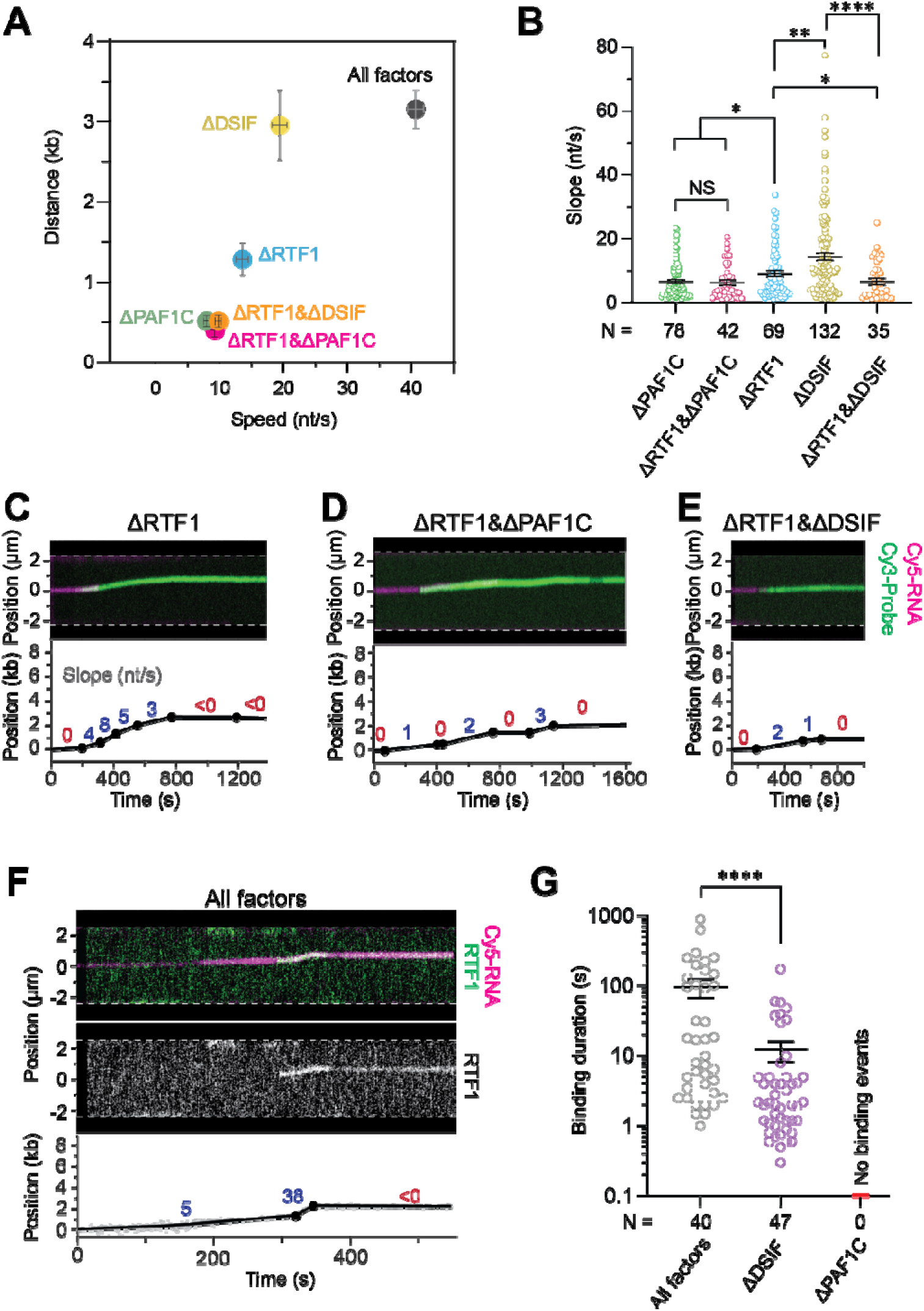
RTF1’s stimulatory effect on EC differentially relies on PAF1C and DSIF. (A) 2D plot showing the average speed and distance for ECs under different conditions (All factors; single omissions: ΔRTF1, ΔPAF1C, ΔDSIF; double omissions: ΔRTF1&ΔPAF1C, ΔRTF1&ΔDSIF). Error bars represent SEM. (B) Dot plot showing the slope of active elongation segments from EC trajectories under ΔPAF1C, ΔRTF1&ΔPAF1C, ΔRTF1, ΔDSIF, and ΔRTF1&ΔDSIF conditions. Bars represent mean ± SEM. NS, not significant; \**P* < 0.05; \*\**P* < 0.01; \*\*\*\**P* < 0.0001. *P* values were from unpaired t tests with Welch’s correction. (C) An example EC trajectory under the ΔRTF1 condition. (Top) Kymograph showing the overlay of Cy5-labeled RNA (magenta) and Cy3-labeled complementary DNA probes (green) which were used to track EC progression. (Bottom) Fitted and segmented trajectory with the slope of each segment indicated. (D) An example EC trajectory under the ΔRTF1&ΔPAF1C condition. Details are the same as (C). (E) An example EC trajectory under the ΔRTF1&ΔDSIF condition. Details are the same as (C). (F) An example EC trajectory with Cy3-labeled RTF1 and Cy5-labeled RNA when the full set of elongation factors were present (All factors). (Top) Kymograph showing the overlay of RTF1 (green) and RNA (magenta) signals. (Middle) The same kymograph but with only RTF1 signals shown. (Bottom) Fitted and segmented trajectory with the slope of each segment indicated. (G) Dot plot showing the lifetime of RTF1 binding events on EC under different conditions. Bars represent mean ± SEM. \*\*\*\**P* < 0.0001. *P* values were from unpaired t tests with Welch’s correction.

Finally, we attached a Cy3 fluorophore to the C-terminus of RTF1 to directly visualize its behavior during elongation. We observed that the appearance of RTF1 signals on EC correlated with transition of the EC from low speed to high speed (Figures 6F and S10D), confirming RTF1’s direct stimulatory effect. We also noticed that RTF1 can bind to the DNA template, which may help RTF1’s search for the EC (Figure S10D). Strikingly, omitting PAF1C completely abolished RTF1 binding events on either EC or DNA (Figure 6G), confirming that PAF1C’s recruitment to EC precedes RTF1. On the other hand, RTF1 was still observed to bind EC in the absence of DSIF, even though the average residence time was significantly reduced (Figure 6G). This is consistent with the double-omission results above, together suggesting that RTF1 can positively regulate EC activity independently of DSIF, but the full activation of EC still requires both factors.

## Discussion

### Mammalian EC is a multi-gear supramolecular machine

The work presented here, for the first time, characterized the basic kinetic properties of mammalian transcription elongation such as its speed, processivity, and pausing frequency in a fully reconstituted system. Our results show that the mammalian EC displays a remarkable level of heterogeneity and dynamic range in its speed, which likely reflects its need to drastically shift gears during the transcription cycle: slow during the early stage prior to the assembly of a complete EC, fast after the escape from promoter-proximal pausing, and then slow again when reaching the 3’ end of genes.^2,6,8^ We further delineated how the acceleration and deceleration of Pol II is controlled by the binding and dissociation of specific factors (Figure 7). Although the factors studied here have all been categorically considered as positive elongation factors, our experiments revealed that they each regulate the elongation speed and distance in a unique and hierarchical manner. The fact that the absence of any single factor (P-TEFb, DSIF, PAF1C, SPT6, RTF1) results in a reduced speed suggests that these factors are not functionally redundant but instead collaborate to meet the demands of the complex cellular environment in multicellular organisms.

**Figure 7.**
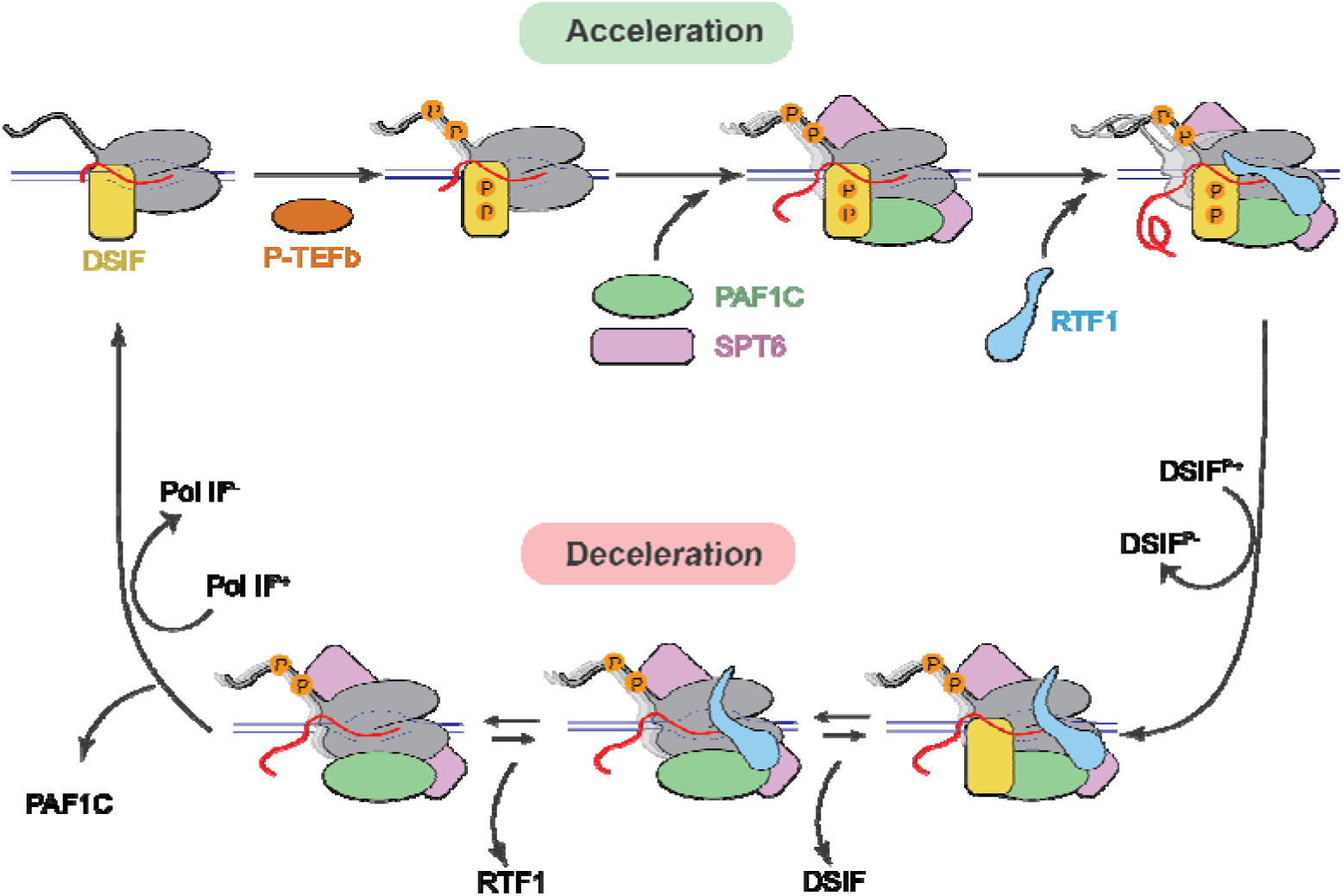
Working model for the kinetic control of the mammalian EC. Each state is defined by a distinct set of associated factors and the phosphorylation state of the EC. P-TEFb poises the EC for active elongation by phosphorylating Pol II and DSIF. DSIF play a dual role in elongation by both enhancing speed and promoting pausing. The recruitment of PAF1C to the EC, stabilized by SPT6, directly speeds up elongation. The final boost in elongation speed is provided by RTF1, whose binding to EC strictly requires PAF1C but not DSIF.

### Critical phosphorylation substrates within the EC

By manipulating the phosphorylation status of specific components of the EC, we showed that the regulation of the elongation speed by P-TEFb is mainly manifested in its phosphorylation of Pol II and DSIF. On the other hand, the phosphorylation of the other CDK9 substrates, including PAF1C and SPT6, does not directly impact the EC speed. The phosphorylation of Pol II CTD is a prerequisite for PAF1C recruitment, which we found to be the most critical event for EC acceleration. The phosphorylation of DSIF greatly stabilizes its own association with EC and possibly mediates RTF1’s stimulatory function.^11^ As such, dephosphorylation of DSIF and Pol II CTD by protein phosphatases (PP1- and PP2A-associated complexes)^48^ may induce the dissociation of speed-stimulating factors and thus the deceleration of EC (Figure 7), potentially facilitating termination or premature termination.

### The multifaceted roles of DSIF

DSIF, specifically its SPT5 subunit, is the only universally conserved transcriptional regulator across domains of life.^31^ We showed that DSIF enhances the elongation speed of the EC and also promotes its pausing, providing a mechanistic basis for its perplexing role in both stimulating and repressing the transcriptional activity reported before.^13,29,35,49,50^ The speed-enhancing effect of DSIF resembles its bacterial homolog NusG, which shares an NGN and KOW domain with SPT5.^51^ Mammalian SPT5 has also evolved additional KOW domains and the C-terminal repeat (CTR), which harbor many phosphorylation sites. We found that the unphosphorylated DSIF loses the speed-enhancing function but still retains the pause-stabilizing function. As such, the exact regulatory function of DSIF can be tuned by its phosphorylation status, making it a versatile elongation factor.

### The PAF1C-RTF1-DSIF nexus

Among all the factors examined in this work, PAF1C exerts the most direct impact on Pol II speed—its binding immediately accelerates the EC. This type of insight highlights the power of real-time single-molecule visualization in unveiling causal effects. Moreover, our data show that RTF1 provides an additional boost to the elongation speed and that it can bind Pol II without DSIF; nevertheless, DSIF significantly stabilizes RTF1’s association with EC. In the EC structure that contains both RTF1 and DSIF, the latch domain of RTF1 makes contacts with the bridge helix of Pol II, which are important for robust EC activity.^11^ We postulate that these contacts are allosterically enabled by RTF1-DSIF interaction, which could explain the observed synergy between these two factors.

Rtf1 is a constitutive component of Paf1C in yeast but becomes a separate entity in metazoans.^52^ Previous results from the *S. cerevisiae* system showed that Spt4/5 is essential for Paf1C recruitment via Rtf1.^53^ In contrast, our findings from the mammalian system reveal that the recruitment of PAF1C to EC only relies on P-TEFb and SPT6 but not on RTF1 or DSIF. Thus, the PAF1C-RTF1-DSIF nexus has acquired new functionalities in higher eukaryotes, allowing more sophisticated kinetic control.

## Conclusion

The elongation speeds we measured under the in vitro reconstituted setting mimic in vivo measurements,^54,55^ and our factor-omission results largely recapitulate the effects observed from in vivo depletion experiments.^12,14,36,38,49^ These agreements support the utility of our single-molecule platform in dissecting the precise functions of individual elongation factors as well as their interplay, and indicate that cells possess efficient mechanisms to overcome the nucleosome barrier during transcription elongation.^56,57^ This study sets the stage for elucidating these mechanisms by visualizing transcription on chromatin templates.

## Acknowledgments

We thank Fei Chen and Gabriella Chua for critical readings of the manuscript. S.L. is supported by the Alfred P. Sloan Foundation, and the Robertson Foundation. Y.X. and X.C. are supported by grants from the National Key R&D Program of China (2021YFA1300100), the National Natural Science Foundation of China (32271242), the XPLORER Prize, and the New Cornerstone Investigator Program.

## Author contributions

Y.W., X.C., Y.X. and S.L. designed the experiments. Y.W. performed the experiments and conducted data analysis. X.C. prepared the protein reagents. M.K assisted with single-molecule data collection and performed structural modeling. J.W.W. wrote data acquisition scripts. J.E.C. developed data analysis software. S.L., Y.W. and Y.X. wrote the manuscript with inputs from all authors.

## Materials and Methods

### Protein purification

#### Pol II

Pol II was isolated from *Sus scrofa* thymus and purified following previously established protocol.^58,59^ The purified Pol II was aliquoted and stored in liquid nitrogen.

#### TFIIS

The full-length ORF of human TFIIS was cloned into a modified pRSFDuet-1 vector and expressed in *E. coli* BL21 (DE3) cells. Cells were grown at 37 °C until reaching an optical density at 600 nm (OD_600_) of around 0.6. The temperature was decreased to 16 °C and protein expression was induced by adding 0.1 mM Isopropyl β-D-1-thiogalactopyranoside (IPTG). Cells were grown for an additional 16 h at 16 °C and were harvested and re-suspended in a buffer containing 30 mM Tris-HCl 8.0, 300 mM NaCl and 20 mM imidazole. Cells were lysed by high pressure at 4 °C for 20 min and the lysate was cleared by centrifugation at 15,000 rpm for 40 min at 4 °C. The supernatant was incubated with Ni-NTA resin for 1 h. Beads were collected and washed with buffer containing 30 mM Tris-HCl pH 8.0, 300 mM NaCl, 25 mM imidazole and eluted with a buffer containing 30 mM Tris-HCl 8.0, 300 mM NaCl and 250 mM imidazole. The elute was dialyzed against a buffer containing 30 mM Tris-HCl pH 8.0 and 300 mM NaCl and supplied with HRV 3C protease to cleave the 6x histidine-SUMO-tag overnight. The eluate after dialysis was loaded onto a Mono S 5/50 GL column (Cytiva), pre-equilibrated with buffer A (30 mM Bis-Tris pH 6.4 and 30 mM NaCl). The target protein was eluted with a gradient from 0% to 100% buffer B (30 mM Bis-Tris pH 6.4 and 1 M NaCl) over 30 column volumes. The peak fractions were pooled, concentrated and dialyzed overnight against a buffer containing 30 mM HEPES pH 7.9, 100 mM KCl, 6 mM MgCl_2_, 2 mM DTT and 5% (v/v) glycerol. The protein was aliquoted, flash-frozen in liquid nitrogen, and stored at −80 °C.

#### P-TEFb

The two full-length ORFs of human P-TEFb were cloned into a modified pCAG vector, and CCNT1 was tagged with an N-terminal protein A tag. The plasmid was transfected into Expi293F cells for overexpression. After cultured at 37 °C for 72 h, cells were harvested and lysed in Lysis buffer containing 30 mM HEPES pH 8.0, 300 mM NaCl, 0.25% CHAPS, 5 mM ATP, 5 mM MgCl_2_, 0.2 mM EDTA, 2 mM DTT, 10% glycerol (v/v), 1 mM PMSF, 1 μg/mL Aprotinin, 1 μg/mL Pepstatin and 1 μg/mL Leupeptin at 4 °C for 30 min. The lysate was clarified by centrifugation at 15,000 rpm for 40 min at 4 °C. The supernatant was incubated with IgG resin for 1.5 h followed by on-column digestion by ULP1 protease at 4°C for 1 h. The immobilized proteins were eluted and further purified by a mono S 5/50 GL column (Cytiva) pre-equilibrated in buffer A (30 mM HEPES pH 7.0, 30 mM NaCl, 0.05% CHAPS, 2 mM DTT, 2 mM MgCl_2_ and 5% (v/v) glycerol). The target protein was eluted with a gradient from 0% to 100% buffer B (same as buffer A except with 1 M NaCl) over 30 column volumes. The peak fractions were pooled, aliquoted, flash-frozen in liquid nitrogen, and stored at −80 °C.

#### PAF1C

The five full-length ORFs of human PAF1C were cloned into a modified pCAG vector. CDC73 was tagged with an N-terminal protein A-SUMO tag, and CTR9 was tagged with an N-terminal protein A-SUMO tag or an N-terminal protein A-SUMO-S6 tag (S6: GDSLSWLLRLLN).^60^ The plasmids were transfected into Expi293F cells for overexpression. After cultured at 37 °C for 72 h, cells were harvested and lysed in Lysis buffer at 4 °C for 30 min. The lysate was clarified by centrifugation at 15,000 rpm for 40 min at 4 °C. The supernatant was incubated with IgG resin for 1.5 h followed by on-column digestion by ULP1 protease at 4 °C for 1 h. The immobilized proteins were eluted and further purified by a mono S 5/50 GL column (Cytiva) pre-equilibrated in buffer A (30 mM HEPES pH 8.0, 30 mM NaCl, 2 mM DTT, 2 mM MgCl_2_ and 5% (v/v) glycerol). The target protein was eluted with a gradient from 0% to 100% buffer B (same as buffer A except with 1 M NaCl) over 30 column volumes. The peak fractions were pooled, aliquoted, flash-frozen in liquid nitrogen, and stored at −80 °C.

#### ELOF1

The full-length ORF of human ELOF1 was cloned into a modified pRSFDuet-1 vector and expressed in *E. coli* BL21 (DE3) cells. Cells were grown at 37 °C until reaching an OD_600_ of around 0.6. The temperature was decreased to 16 °C and protein expression was induced by adding 0.1 mM IPTG. Cells were grown for an additional 16 h at 16 °C, and harvested and re-suspended in buffer C containing 50 mM Tris-HCl 8.5, 700 mM NaCl, 0.7 mM β-mercaptoethanol, 5% (v/v) glycerol and 25 mM imidazole. Cells were lysed by high pressure at 4 °C for 20 min and the lysate was cleared by centrifugation at 15,000 rpm for 40 min at 4 °C. The supernatant was incubated with Ni-NTA resin for 1 h. Beads were collected and washed with buffer C. The bound protein was eluted using a buffer containing 30 mM Tris-HCl pH 8.0, 300 mM NaCl and 250 mM imidazole, followed by overnight treatment with ULP1 protease. The elute was further purified by a mono Q 5/50 GL column (Cytiva) pre-equilibrated in buffer A (30 mM Tris-HCl 8.5, 50 mM NaCl, 2 mM DTT and 5% (v/v) glycerol). The target protein was eluted with a gradient from 0% to 100% buffer B (same as buffer A except with 1 M NaCl) over 30 column volumes. The peak fractions were pooled, aliquoted, flash-frozen in liquid nitrogen, and stored at −80 °C.

#### SPT6

The full-length ORF of human SPT6 was cloned into a modified pCAG vector containing an N-terminal protein A tag. The plasmid was transfected to Expi293 cells for overexpression. After cultured at 37 °C for 72 h, cells were harvested and lysed in a buffer containing 30 mM HEPES pH 8.0, 300 mM NaCl, 0.25% CHAPS, 5 mM ATP, 5 mM MgCl_2_, 0.2 mM EDTA, 2 mM DTT, 10% (v/v) glycerol, 1 mM PMSF, 1 μg/mL Aprotinin, 1 μg/mL Pepstatin and 1 μg/mL Leupeptin at 4 °C for 30 min. The lysate was clarified by centrifugation at 15,000 rpm for 40 min at 4 °C, and the supernatant was incubated with IgG resin for 1.5 h followed by on-column digestion by ULP1 protease at 4 °C for 1 h. The immobilized proteins were eluted and further purified by a mono Q 5/50 GL column (Cytiva) pre-equilibrated in buffer A (30 mM HEPES 8.0, 30 mM NaCl, 2 mM DTT, 2 mM MgCl_2_ and 5% (v/v) glycerol). The target protein was eluted with a gradient from 0% to 100% buffer B (same as buffer A except with 1 M NaCl) over 30 column volumes. The peak fractions were pooled, aliquoted, flash-frozen in liquid nitrogen, and stored at −80 °C.

#### DSIF, RTF1, IWS1

These proteins were purified in a similar way to that described for SPT6. The two full-length ORFs of human DSIF were cloned into a modified pCAG vector, and SPT4 was tagged with an N-terminal protein A-SUMO tag or an N-terminal protein A-SUMO-spy tag (spytag3: RGVPHIVMVDAYKRYK).^61^ The plasmids were co-transfected into Expi293F cells for overexpression. The ORF of full-length human RTF1 with a C-terminal spy tag or truncated RTF1 (residues 126-710) was cloned into a modified pCAG containing an N-terminal protein A tag and overexpressed in Expi293F cells. The truncated RTF1 showed a similar activity of stimulating transcription as the full-length RTF1.^11^ The full-length ORF of human IWS1 was cloned into a modified pCAG vector containing an N-terminal protein A tag and overexpressed in Expi293F cells. After cell lysis, the lysate was applied to an IgG resin followed by on-column digestion. The eluate was further purified by a mono Q 5/50 GL column (Cytiva). The peak fractions were pooled, aliquoted, snap frozen and stored at −80°C.

### Protein labeling

To site-specifically label the PAF1C complex, the S6-PAF1C, Sfp synthase and Cy3-CoA dye (SiChem) were incubated at a 1:20:100 molar ratio for 4 h at room temperature and overnight at 4 °C in the presence of 10 mM MgCl_2_. Excess dye and Sfp synthase were removed by a 50-kDa Amicon spin filter (Millipore) with a buffer containing 30 mM HEPES pH 7.9, 250 mM NaCl, 5 mM MgCl_2_, 5 mM DTT and 5% (v/v) glycerol. The labeled protein was evaluated via SDS-PAGE gel electrophoresis. The gel was scanned by Typhoon FLA 7000 (GE Healthcare). The labeling efficiency was estimated to be 92%. Final protein samples were aliquoted, flash frozen, and stored at −80 °C.

DSIF and RTF1 were labeled through the conjugation between fluorescently labeled spycatcher3 protein and the spy tag.^61^ Spycatcher3 with a single cysteine residue (Bio-Rad) was first incubated with 10 mM tris(2-carboxyethyl)phosphine (TCEP) (Sigma-Aldrich) at room temperature for 1 h to reduce the thiol group. Cy3 maleimide mono-reactive dye (Cytiva) was then added with a 1:5 protein-to-dye molar ratio. The reaction was conducted at room temperature for 2 h and overnight at 4 °C. Excess dye and TCEP were removed by running Bio-Spin-6 column (Bio-Rad) twice. The labeling efficiency for Cy3-spycatcher3 was estimated to be 141%. Spy-tagged DSIF or full-length RTF1 was mixed with Cy3-spycatcher3 at room temperature for 20 min, with the optimal molar ratio pre-determined through titration.

### DNA substrate preparation

To generate the DNA template for single-molecule experiments, an 8-kb plasmid ply-8 was adapted from a 13.8-kb plasmid plw83 (ref. ^62^) where a 6-kb fragment of plw83 containing multiple BbsI recognition sites was truncated. One new single BbsI recognition site was inserted into the truncated plw83 plasmid. A cassette harboring seven tandem repeats of a 21-bp sequence (5’-AGACACCACAGACCACACACA) was placed 76 bp downstream of the BbsI site. Ply-8 was digested using BbsI to generate the linear DNA template (Step 1 in Figure S2A). The linearized DNA was ethanol precipitated at -20 °C for 1 h by 3 volumes of cold ethanol and 300 mM NaOAc pH 5.2. Precipitated DNA was recovered by centrifugation at >20,000 g at 4 °C for 30 min. The DNA pellet was washed by 75% ethanol, air-dried, and resuspended in ddH_2_O. To increase the ligation efficiency and specificity, the (5’CCCA) 4-nt ssDNA overhang in the linearized DNA was extended to 15 nt by ligation with bridge and overhang oligonucleotides (Step 2 in Figure S2A). Specifically, the purified linearized ply-8 was mixed with Overhang-oligo (5’Phos-TGGGTGGTGTTCCGCACATGCTGAGTCGAGCTTA) (Integrated DNA Technologies) and Bridge-oligo (5’Phos-ATGTGCGGAACACCA) (Integrated DNA Technologies) with a molar ratio of 1:100:125 in a buffer containing 5 mM Tris-HCl pH 7.5 and 50 mM NaCl. The mixture was heated to 75 °C for 10 min and then cooled down to room temperature gradually for 1 h. T4 ligation buffer and T4 ligase (final concentration of 2U/μL) was added and incubated at 16 °C overnight. The ligation product was purified twice by DNA size-selective magnetic beads (Sergi Lab Supplies) to remove the excess oligos (following the manufacturer’s protocol) and eluted by ddH_2_O.

The 5’ overhangs of the purified ligation product were filled in with biotinylated nucleotides by the exonuclease-deficient DNA polymerase I Klenow fragment (New England BioLabs) to create terminally biotinylated DNA for bead conjugation (Step 3 in Figure S2A). The fill-in reaction was conducted by incubating 100-200 nM linearized plasmid DNA, 33 μM each of biotin-14-dATP/biotin-14-dCTP (Jena Bioscience), and 10 U of Klenow fragment in 1x NEB2 buffer at room temperature for 2 h. To stop the reaction, EDTA was added at a final concentration of 10 mM and the reaction mixture was heat inactivated at 75 °C for 20 min. The DNA was then ethanol precipitated overnight at -20 °C and resuspended in ddH_2_O.

*Template-oligo* (5’Phos-CGTTTGTGTTTTCGGGTCTCCCTCGTTTCTGGCTTGGGTTGGCTTTTC<u>GCCGTGTCG</u>TACATCATACTACTCCTACCAGGAAGCATAAGCTCGACTCAGC, underlined indicates the RNA hybridization region) (Integrated DNA Technologies) or *Nontemplate-oligo* (5’Phos-TGCTTCCTGGTAGGAGTAGTATGATGTACGACACGGCGAAAAGCCAACCCAAGCCAGAAACGAGGGAGACCCGAAAACACAAACGTAAGCTCGACTCAGC) (Integrated DNA Technologies) was ligated to the biotinylated 8kb-DNA from Step 3 to create non-template-strand DNA (8k-NTS-DNA) and template-strand DNA (8k-TS-DNA) handles used for elongation complex assembly (Step 4 in Figure S2A). Specifically, Template-oligo or Nontemplate-oligo was mixed with biotinylated 8kb-DNA with a molar ratio of 5:1 in a buffer containing 5 mM Tris-HCl pH 7.5 and 50 mM NaCl. After annealing as described in Step 2, T4 ligation buffer and T4 ligase (final concentration of 2U/μL) was added and ligated at 16 °C overnight. The ligation product was purified once by DNA size-selective magnetic beads and eluted by ddH_2_O with the final DNA concentration ∼1300 ng/μL (or 250 nM). The final products were aliquoted and stored in -80 °C.

### Single-molecule experiments

#### Formation of elongation complexes

To prepare elongation complexes as shown in Figure S2B, 2 μL of 8k-TS-DNA (250 nM) was mixed with 2 μL of EB40 buffer (20 mM Tris-HCl pH 8.0, 40 mM KCl and 5 mM MgCl_2_) and 2 μL of Cy5-RNA-primer (/5ATTO647NN/UUUUUUUUUUUAUAU<u>CGACACGGC</u>, underlined indicates the hybridization region; 10 μM, dissolved in EB40) (Integrated DNA Technologies). The mixture was subjected to an annealing procedure: 55 °C for 30 s, 50 °C for 30 s, 45 °C for 30 s, 40 °C for 2 min, 37.5 °C for 2 min, 35 °C for 2 min, 30 °C for 2 min, 27 °C for 2 min, 25°C for 10 min, and 4 °C indefinitely. Following annealing, 0.5 μL of freshly prepared 0.1 M DTT, 0.5 μL of RNase Inhibitor murine (40 U/μL) and 2 μL of Pol II (3 μM) were added to the DNA-RNA hybrid solution. The mixture was incubated at 30 °C for 8 min, followed by 37 °C for 2 min, to assemble the transcription bubble. 2 μL of 8k-NTS-DNA (250 nM), 0.52 μL of ATP (25 mM) and 2 μL of P-TEFb (4 μM) were added and incubated at 30 °C for 20 min. Finally, the nicks from 8k-TS-DNA/8k-NTS-DNA hybridization were sealed by adding 0.75 μL of T4 ligase (2000 U/μL) and incubating at room temperature for 1 h. The assembled elongation complex was ready to be loaded into the microfluidic chip for single-molecule assays.

#### Single-molecule data collection

Single-molecule experiments were performed at room temperature on a LUMICKS C-Trap instrument. Channels of the microfluidic chip were passivated by first flowing BSA (0.1% (w/v) in 1x PBS, Sigma-Aldrich) and then Pluronic F127 (0.5% (w/v) in 1x PBS) for 10 min each. Streptavidin-coated polystyrene beads (2.14-μm diameter, Spherotech, diluted in 1x PBS), 16-kb DNA-EC (100 pM in imaging buffer), imaging buffer, and transcription buffer (all four ribonucleotides at 1 mM each and elongation factors in imaging buffer) were injected into channels 1-4, respectively. Imaging buffer included an oxygen scavenging system (10 nM protocatechuate-3,4-dioxygenase (Sigma-Aldrich) and 2.5 mM protocatechuic acid (Sigma-Aldrich)), 0.3 U/μL RNase inhibitor murine, 5 mM freshly made DTT and a triplet-state quenching cocktail (1 mM cyclooctatetraene, 1 mM 4-nitrobenzyl alcohol, and 1 mM Trolox (Sigma-Aldrich)). The final concentrations of the elongation factors in transcription buffer for the All factors condition were: PAF1C 50 nM, SPT6 120 nM, RTF1_126-710_ 200 nM, TFIIS 0.5 μM, DSIF 200 nM, IWS1 60 nM, ELOF1 1 μM, and P-TEFb 50 nM. In those experiments that involved fluorescently labeled factors, a lower concentration was used to reduce the fluorescence background (Cy3-DSIF^P+^ 12.5 nM, Cy3-DSIF^P-^ 12.5 nM, Cy3-PAF1C 25 nM, Cy3-RTF1_FL_ 33 nM). To prepare the transcription buffer in channel 4, a pre-phosphorylation step was conducted before mixing the elongation factors with the imaging buffer. Specifically, 3-fold concentrated elongation factors were incubated with 1 mM ATP at 30 °C for 30 min to allow P-TEFb-mediated phosphorylation of a specific set of factors. For the DSIF^P-^ condition, DSIF was omitted from this step and subsequently added to the transcription mix together with 200 nM CDK9 inhibitor flavopiridol (Selleck Chemicals LLC). For the Pol II^P-^ condition, P-TEFb was omitted during the steps of EC formation (Figure S2B) and 200 nM flavopiridol was included in the transcription buffer to prevent phosphorylation of Pol II in channel 4. For the DSIF^P+^&Pol II^P+^ condition, only DSIF was incubated with P-TEFb and ATP, and flavopiridol was included in the transcription buffer.

A single DNA tether was caught between a pair of streptavidin beads held in optical traps, and the tether was extended at 5 pN of tension. The tether was moved to channel 3 for confocal scanning using a 638-nm laser to verify the existence of assembled EC. Empty tethers and those with protein aggregates were discarded. Kymographs were generated using the Bluelake software (LUMICKS) via confocal line scanning through the center of the two beads at 0.2 s/line (pixel time: 0.2 ms). Data acquisition started in channel 3 for 20 s for baseline calibration and continued in channel 4. When applicable, 10 nM of Cy3-labeled DNA oligo probes (/5Cy3/AGTGTGTGGTCTGTGGTGTCT, Integrated DNA Technologies) were included in channel 4 to detect nascent RNA.

### Single-molecule data analysis

#### Data acquisition

Single-molecule force and fluorescence data from the .h5 files generated by Bluelake were processed using the *lumicks.pylake* Python library, supplemented by other Python modules integrated into a custom GUI script, *CTrapVis.py* (http://harbor.lumicks.com/single-script/c5b103a4-0804-4b06-95d3-20a08d65768f). This script was used to export confocal scans and kymographs in TIFF format, as well as to extract real-time data for force, distance, and photon counts. The positions of Cy5-RNA and Cy3-DNA probes on the template over time were determined from kymographs using a custom script, *kymotracker_calling_script.py* (https://harbor.lumicks.com/single-script/4db9d63e-1f93-0c0-9b90-e99066469578), which implemented the greedy line-tracking algorithms provided by Python package *lumicks.pylake* to define line traces. Optimal tracking of EC was achieved by setting a pixel threshold of 1 photon count for Cy5 signals and 2 photon counts for Cy3 signals, with a line width of 12 pixels, a window of 35 frames, and a minimum line length of 20 frames. The extracted trajectories were aligned based on the starting position of the Cy5-RNA using data collected in channel 3. Distance traveled in nanometers was converted to base pairs using a scaling factor of 0.29 nm/bp, reflecting the extension of B-form DNA under 5 pN of tension.

#### Segmentation analysis of extracted transcription trajectories

##### Structure of the data

A condition is specified by the chemical composition of the elongation complex. Each elongation complex contains some subset of the 8 human elongation factors. Here we have *C* different conditions corresponding to C different elongation complexes. We refer to each condition by the variable *c* = 1, 2, … , *C*; there are 14 conditions in total, so C=14. For example, the elongation complex with all factors is *c* = 1; the elongation complex without TFIIS is *c* = 2; the elongation complex without IWS1 is *c* = 3, and so on. In an Excel data file, each condition has its own sheet.

For each elongation complex *c*, we have T(c) trajectories. Different conditions c may have different numbers of trajectories T(c). In condition c, each trajectory t = 1, 2, …, T(c) consists of a pair of column vectors (S(c,t), K(c,t)) which are of the same length L(c,t). The length is measured by the number of elements (that is, the number of times an observation was recorded) for trajectory t in condition c in S(c,t) and also in K(c,t). These vectors appear in adjacent vertical columns in an Excel spreadsheet. The first row of each sheet gives a unique alphanumeric identifier for each trajectory. S is a mnemonic for “seconds.” K is a mnemonic for “kilobases.” For trajectory t in condition c, the jth element of S(c,t), which we label S(c,t,j), is the time at which the jth observation is recorded. Time is measured in seconds from the beginning of a trajectory. The corresponding jth element K(c,t,j) of K(c,t) is the transcription length at time S(c,t,j). Transcription length is measured in kilobases.

Each trajectory is segmented separately. We simplify the notation used to describe how a trajectory is segmented: assume a fixed condition c and a fixed trajectory t in condition c. We describe the segmentation analysis as applied to one pair of vectors (S, K) each of length L. For j = 1, 2, …, L, the j^th^ time of observation is S(j) and the transcription length at time S(j) is K(j). The goal of the following analysis is to approximate the plot of K as a function of S by a sequence of linear segments (Figures 1C and S3) which may have slope 0 (horizontal segment) or may have a positive slope (as transcription proceeds) or a negative slope (as transcription is reversed or backtracking of Pol II). Each segment is separated from its preceding segment (if any) and from its following segment (if any) by a change point, a time at which the estimated slope changes.

##### MATLAB software

Our code is written in Matlab (Mathworks R2023b). The user chooses several parameters that govern the software. These parameters are explained further below.

1. *readmat* = 0 means read the data from an Excel spreadsheet; readmat = 1 means read data from a .mat file where it was stored previously;
2. *Nsheets* = *C* is the number of conditions;
3. *globalmaxchangepoints* = 15 limits maximum number of changepoints (NCP);
4. *mintimegap* = 3 seconds sets minimal time (s) between changepoints; if two changepoints are initially estimated within ’mintimegap’, they are replaced by one changepoint at the midpoint;
5. *δ* = 10^-6^, the width of the interval of random perturbations of the recorded times, to break ties.

When the data are read from an Excel file, the user must specify which columns house data. For each condition (Excel sheet) c, our software calculates the number T(c) of trajectories from the number of columns that house data. When the data are read from a .mat file, the saved variables are:

1. *Nsheets* = C, the number of sheets or conditions;
2. *sheetnames*, a list of alphanumeric names for each condition;
3. *Ntrajectories* = T = (T(1), … ,T(C)), a vector of the number T(c) of trajectories in each condition c, for c = 1, 2, …, C;
4. *outnum*, all the numerical values in each sheet;
5. *outtxt*, all the alphanumeric column headings, which are the names of the individual trajectories;
6. *sheetnames*, *stringsheetnames*, two formats for the names of the sheets or conditions;
7. time = S(c,t,j), a cell array of the time observations in condition c for trajectory t for j = 1, …, L(c,t), where L(c,t) is the number of observations in condition c for trajectory t;
8. bp = K(c,t,j), a cell array of the observations of transcription length at each time in condition c for trajectory t for j = 1, …, L(c,t), where L(c,t) is the number of observations in condition c for trajectory t;
9. *trajectorynames*, a cell array of length c (one cell for each condition) of the names of the individual trajectories, taken from *outtxt*; the j^th^ cell contains a vector of length T(c), each element of which is a string with the name of the corresponding trajectory.

Sometimes the identical reported time will occur more than once. To break all such ties so that the observations occur in a unique order, we add to each time an independently chosen small random quantity uniformly distributed between -*δ* and +*δ*. We set *δ* = 10^-6^ seconds; the user can change the value at will. After this random perturbation, the vectors S and K are sorted in increasing order of the (perturbed) time element in S.

##### Bayesian estimation of the number of changepoints

We used the Matlab version of *Rbeast*. *Rbeast* is available in Matlab, R, Python, and C. The source code and installation instructions are available at https://github.com/zhaokg/Rbeast. According to the website, “BEAST (Bayesian Estimator of Abrupt change, Seasonality, and Trend) is a fast, generic Bayesian model averaging algorithm to decompose time series or 1D [one-dimensional] sequential data into individual components, such as abrupt changes, trends, and periodic/seasonal variations.^63^ BEAST is useful for changepoint detection (e.g., breakpoints, structural breaks, joinpoints, regime shifts, or anomalies), trend analysis, time series decomposition (e.g., trend vs. seasonality), time series segmentation, and interrupted time series analysis.” The specific version we used is *beast_irreg* with parameters ‘deltat’,1,’season’,’none’.

Rbeast operates by estimating a probability (called *q*(1)) for the possibility that there is just one change point (without specifying the location of that change point). Then, conditional on one change point, Rbeast estimates a probability (called *q*(2) < *q*(1)) for the possibility that there is just one more change point. Then, conditional on two change points, Rbeast estimates a probability (called *q*(3) < *q*(2)) for the possibility that there is just one more change point (i.e., a total of three); and so on, possibly up to our chosen *globalmaxchangepoints*. We find the maximal number of change points, called *MaxCP*, such that the cumulative sum of the first *MaxCP* elements of *q* does not exceed 0.99.

##### Locating the change points for linear segments

With this number *MaxCP* of change points, we identify the times at which those change points occur using the Matlab function *ischange* for K as a piecewise (not necessarily continuous) linear function of the perturbed times (using the parameter ‘linear’) and limiting the maximum number of changes to *MaxCP* (using ’MaxNumChanges’,MaxCP). The *ischange* algorithm is based on Killick et al. ^64^ With the parameter ‘linear’, *ischange* fits straight lines to each segment between consecutive change points but the successive straight lines may not connect into one continuous trajectory. If the difference in seconds between two change points is less than our chosen value for the parameter *mintimegap* (3 s), then we replace the times at which both of those change points occur by a single time located at the arithmetic average of the original two times. This adjustment excludes any segments of duration less than *mintimegap*. We record the number of change points before and after this adjustment to the data as *NchangepointsBefore* and *NchangepointsAfter*, respectively.

##### Finding a continuous piecewise linear approximation

Given the (perturbed) times S and corresponding kilobase pairs transcribed K, and given the locations of the change points, we fit a continuous piecewise linear approximation to K using *fitPiecewiseLinearFunctionGolovchenko.m* written by Nikolai Golovchenko (2004) (https://www.golovchenko.org/home/pwl_fit). This code fits “a piecewise continuous [linear] function f(x) to the pairs of data points (x,y) such that the sum of squares of error is minimal.” It outputs x0, the values of x that define ends of segments of the fitted function f(x), and p, the end points of the linear segments, p = f(x0).

##### Tabulated results

A segment is defined by four numbers: its initial (perturbed) time *S*_O_, its initial kb *K*_O_, its final (perturbed) time *S*_1_ and its final kb *K*_1_. The duration of a segment is defined as *S*_1_ *- S*_O_. The length of a segment is defined as *K*_1_ *- K*_O_. The slope of a segment *BpPerSecBetwCPs* is defined as the ratio of length divided by duration, (*K*_1_ - *K*_O_)/(*S*_1_ - *S*_O_). We define a segment to belong to active transcription (slopeClass 1) if and only if its slope satisfies *BpPerSecBetwCPs* greater than or equal to 1 nt/s and its length exceeds 100 nt. Such a segment has progressing transcription. We define a segment to belong to pause (slopeClass 0) if and only if its slope is less than 1 nt/s. We tabulated the principal characteristics of all the segments of all the trajectories of all the conditions in a table with these columns: sheet (condition), trajectory (or trace) in that condition, segment number in that trajectory, start time for that segment, duration (seconds) of that segment, segment length (kbp), slope, and slopeClass. This table has one row for each segment. Further statistical analysis is based on this table.

#### Analysis of active EC fraction and transcribed length

The parameter “active EC fraction” (Figure S5A) indicates the fraction of EC that displayed transcription activity (emergence of the DNA probe signals) under a given condition. For the All factors condition, the maximal distance that an EC traveled before stalling was averaged and reported as the parameter “transcribed length” (Figure S5B). For all the other conditions (e.g. condition X), the transcribed length was calculated by multiplying the averaged maximal distance by a scaling factor (active EC fraction_X_ / active EC fraction_All factors_).

### Bulk transcription assays

To prepare the EC for bulk assays, 5 μL of *Bulk-template-TS* (0.6 μM, dissolved in EB40 buffer) was mixed with 1.25 μL of Cy5-RNA primer (3 μM, dissolved in EB40 buffer). The mixture was subjected to an annealing procedure: 55 °C for 30 s, 50 °C for 30 s, 45 °C for 30 s, 40 °C for 2 min, 37.5 °C for 2 min, 35 °C for 2 min, 30 °C for 2 min, 27 °C for 2 min, 25 °C for 10 min, and 4 °C indefinitely. Following annealing, 2.5 μL of freshly prepared 0.1 M DTT, 2.5 μL of RNase inhibitor murine (40 U/μL) and 5 μL of Pol II (1 μM) were added to the DNA-RNA hybrid solution. The mixture was incubated at 30 °C for 8 min, followed by 37 °C for 2 min. 5 μL of *Bulk-template-NTS* (0.6 μM, dissolved in EB40) was added and incubated at 30 °C for 10 min. 2.4-μL aliquots of EC were mixed with 7.6 μL of elongation factor mixture (final concentration: 100 nM P-TEFb, 120 nM SPT6, 50 nM PAF1C, 200 nM RTF1_126-710_, 500 nM TFIIS, 200 nM DSIF, 1 μM ELOF1 and 60 nM IWS1) or under conditions where individual factors were omitted (Figure S1B). 1 mM ATP was added to the mixture and incubated for 20 min at 30 °C to allow phosphorylation. All four ribonucleotides were added to 1 mM each to allow transcription for 20 min at 30 °C. 2 μL DNase 10x buffer and 2 μL DNase I (RNase-free) (2U/μL, New England Biolabs) were added and incubated for 10 min at 37 °C to digest the DNA. 8 μL of stop buffer (100 mM Tris-HCl pH 7.5, 150 mM EDTA and 4 M urea) and 1 μL of protease K were added and incubated at 37 °C for 20 min. Finally, the RNA products were mixed with 2x RNA loading buffer and loaded on a 10% Urea-PAGE gel after incubating at 90 °C for 5 min. The gel was scanned by Typhoon FLA 7000 (GE Healthcare) and quantified by ImageJ.

*Bulk-template-TS* (146 nt, Integrated DNA Technologies): 5’CTGCGCGTAATCTGCTGCTTGCTTGCAAACTATCCGGTAACTATCGTCTTGAGTCCT ATGCTTGCCGCTGAGTCGAGCTTAGTCACCATCGATTGTCGCTTGGGTTGGCTTTTCGCCGTGTCGTGGCTGAGCTCTGGCTGTAAGTG (underlined indicates the RNA hybridization region)

*Bulk-template-NTS* (146 nt, Integrated DNA Technologies): 5’CACTTACAGCCAGAGCTCAGCCACGACACGGCGAAAAGCCAACCCAAGCGACAATCGATGGTGACTAAGCTCGACTCAGCGGCAAGCATAGGACTCAAGACGATAGTTACCGGATAGTTTGCAAGCAAGCAGCAGATTACGCGCAG

### In vitro phosphorylation

To phosphorylate Pol II, 1 μL of Pol II (3 μM) was mixed with 1, 2, 3, or 4 μL of P-TEFb (1 μM) in the presence of 1 mM ATP and 2 mM MgCl_2_. To phosphorylate DSIF, 2 μL of DSIF was mixed with 2.5 μL of P-TEFb (1 μM) in the presence of 1 mM ATP and 2 mM MgCl_2_. The reaction was conducted at 30 °C for 30 min. As a negative control, 2 μL of DSIF was mixed with 1 μL of Lambda Protein Phosphatase (New England Biolabs) in the presence of 1x NEBuffer and 0.1 mM MnCl_2_. The reaction was conducted at 30 °C for 30 min. The products were evaluated by SuperSep Phos-tag Precast 7.5% Gels (FUJIFILM Irvine Scientific).

### Structural modeling

To generate a composite structural model that includes all the elongation factors (except P-TEFb) used in our system, we first downloaded the model of human EC containing PAF1C, SPT6, DSIF and RTF1 (PDB: 6TED) using ChimeraX, and then added a human Pol II structure containing TFIIS (PDB: 8A40), a TC-NER structure containing ELOF1 (PDB: 8B3F), and a yeast EC structure containing the IWS1 homolog Spn1 (PDB: 7XN7), superimposing the Pol II in each structure. Residues 550-687 of the AlphaFold 2 model of human IWS1 (AF: Q96ST2) was superimposed to Spn1 in the yeast EC. For AlphaFold 3 predictions of the Pol II-PAF1C-SPT6 interaction, two scenarios were modeled: CDC73^Ras^ + CTD^S2P,S5P^ (binary prediction), and CDC73^Ras^ + CTD^S2P,S5P^ + SPT6^SH2^ + CTD linker (quaternary prediction). The AlphaFold webserver (https://alphafoldserver.com) was used and accessed on 12/01/2024 for the binary prediction and 12/03/2024 for the quaternary prediction. For CDC73^Ras^, residues 351-531 were chosen. For SPT6^SH2^, residues 1323-1520 were chosen. For CTD^S2P,S5P^, we used the sequence (YpSPTpSPSYpSPTpSPSYpSPTpSPS) that was shown to bind yeast Cdc73 (ref.^41^). For the CTD linker, we chose RPB1 residues 1523-1553, which was shown to bind SPT6^SH2^ (ref. ^10^). Predictions were loaded using ChimeraX. The three most highly ranked models for both the binary and quaternary predictions adopt an almost identical trajectory. For better visualization between the two models, the third prediction for both the binary (pTM = 0.66) and quaternary (pTM = 0.78) predictions was chosen. The proposed model in Figure S9C was built using the human EC structure (PDB: 6TED) and the quaternary prediction, with all chains hidden except Pol II, SPT6^Core^ and PAF1C.

---

**Figure S1.**
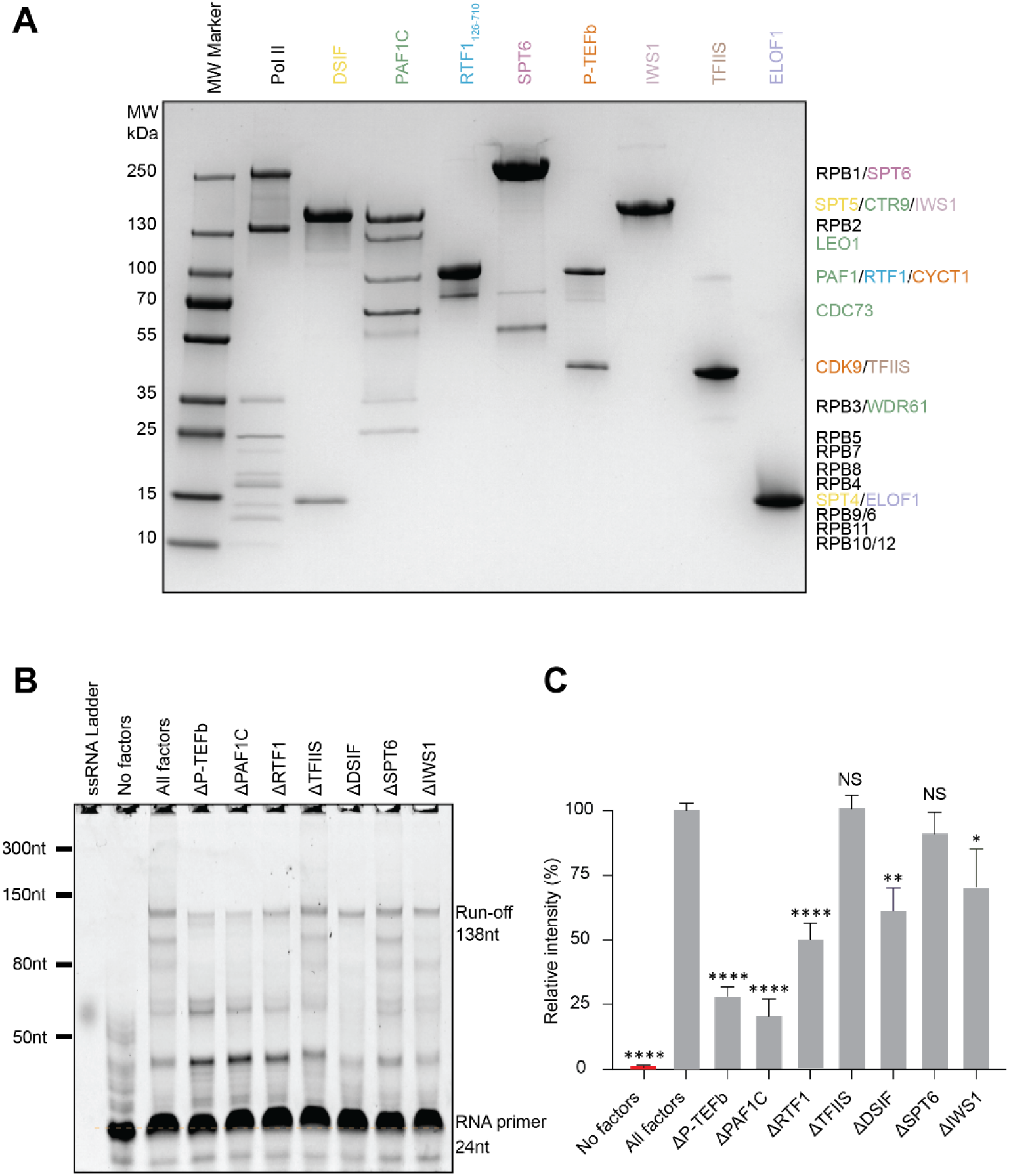
Biochemical evaluations of the proteins used in this study. (A) Coomassie-stained SDS-PAGE gel showing the purified mammalian RNA polymerase II (Pol II) and elongation factors. Protein subunit identities based on their molecular weights are annotated on the right. (B) Bulk transcription assay using the proteins above and a synthetic DNA scaffold with a Cy5-labeled RNA primer. Reconstituted EC (50 nM) was incubated with the full set of elongation factors (All factors) or under conditions where one factor was omitted, in the presence of saturating NTPs (1 mM each). RNA extension was monitored on a denaturing gel by Cy5 fluorescence. The positions of the RNA primer (24 nt) and the run-off product (138 nt) are indicated. Results are representative of three independent replicates. (C) Quantification of the run-off RNA intensity for the same conditions as described in (B) normalized and compared to the All factors condition. NS, not significant; \**P* < 0.05; \*\**P* < 0.01; \*\*\*\**P* < 0.0001. *P* values were from unpaired t tests with Welch’s correction.

**Figure S2.**
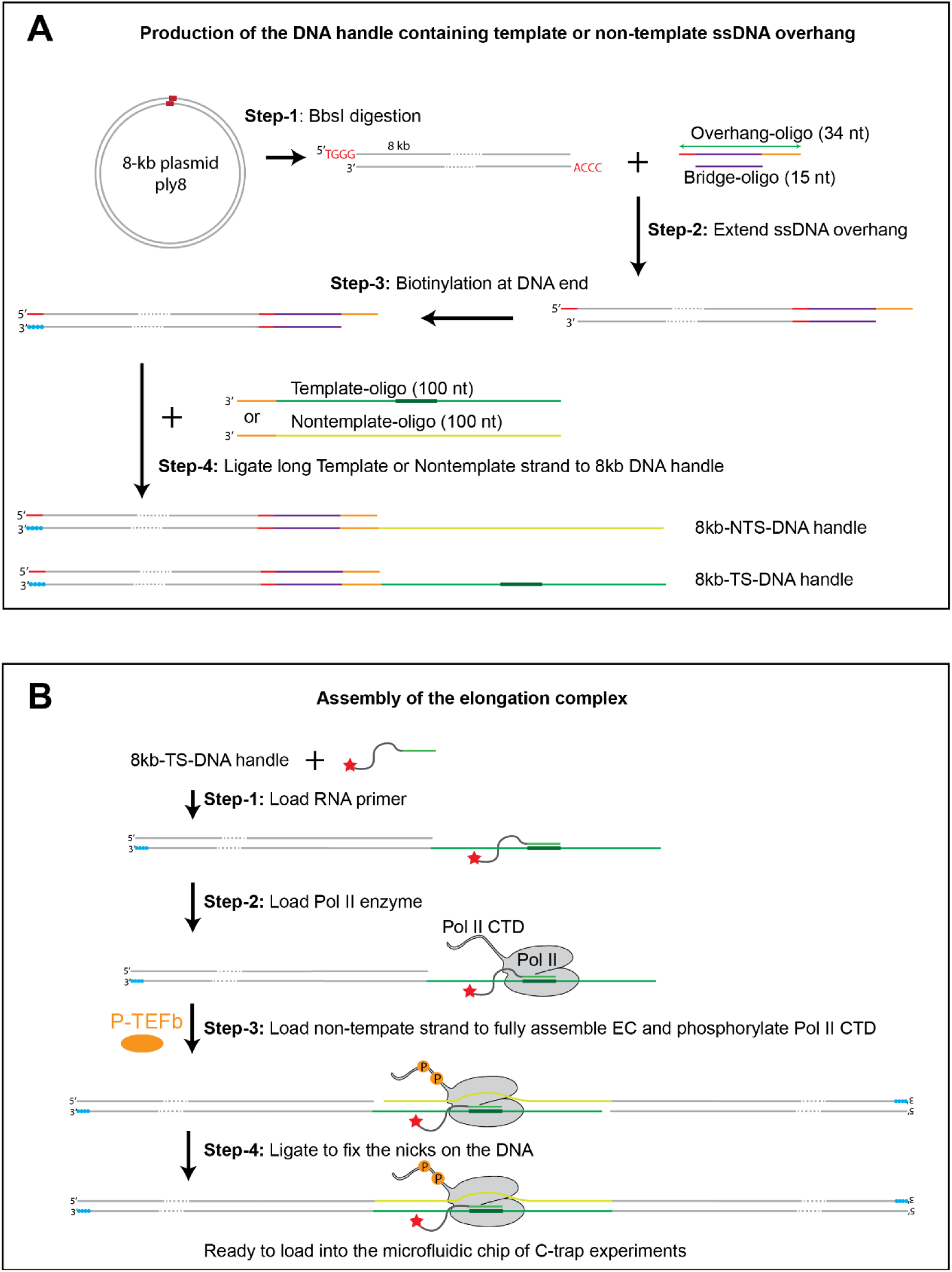
Workflow of EC assembly for single-molecule experiments. (A) Procedure for producing the 8-kb-long non-template-strand (NTS) and template-strand (TS) DNA handles. (B) Procedure for assembling the mammalian EC using the DNA constructs in (A).

**Figure S3.**
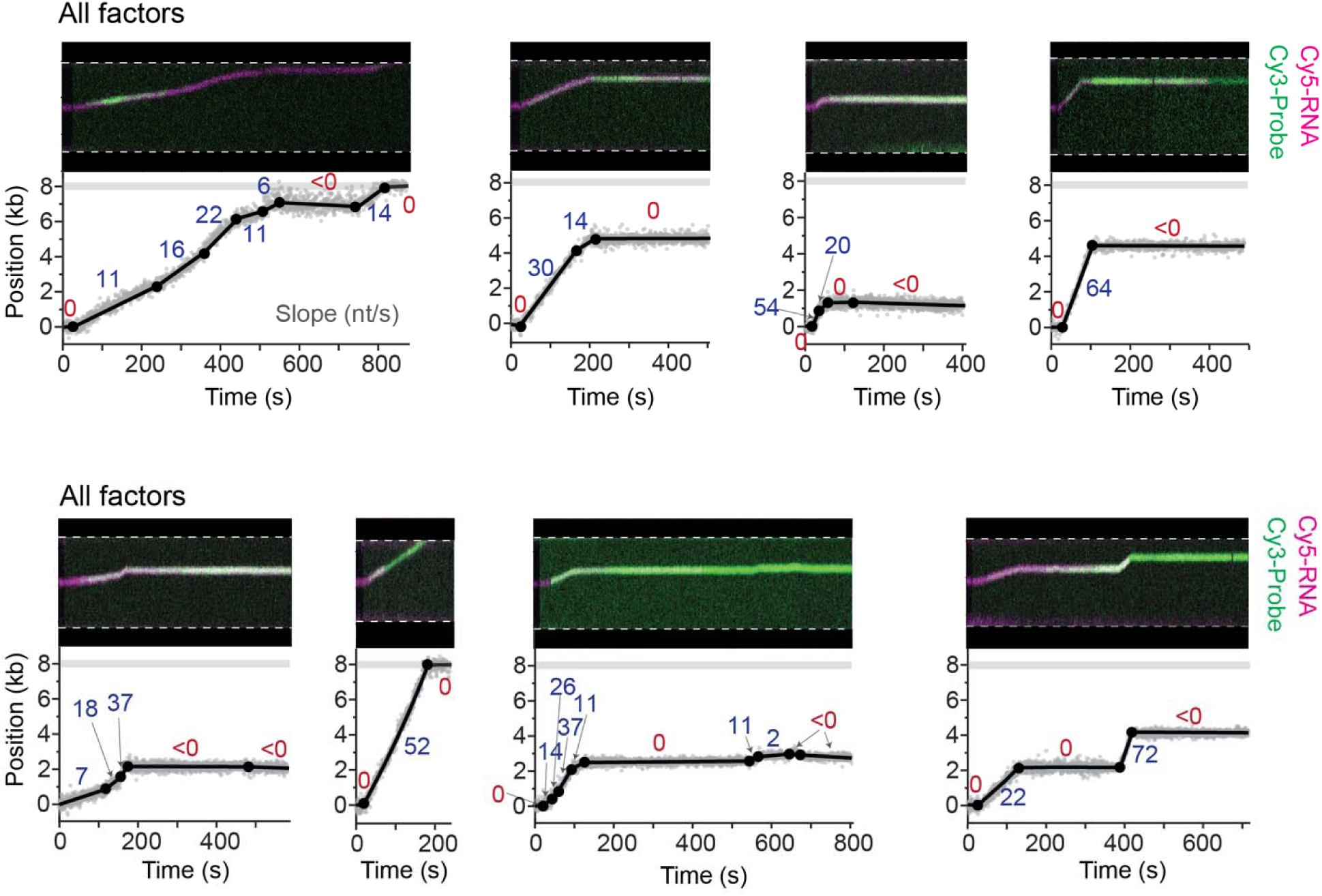
Additional example trajectories of mammalian transcription elongation. In each example, the kymograph (Top) and the corresponding extracted transcription trajectory (Bottom) are shown. Cy5-labeled RNA (magenta) and Cy3-labeled complementary DNA probe (green) in the kymograph were used to track the EC progression. The gray dots represent raw data extracted from the kymograph and the black line indicates the fitted trajectory using a Bayesian framework for changepoint detection. Filled circles denote changepoints, dividing the trajectory into discrete linear segments. The slopes for each segment are indicated. The gray horizontal bar indicates the end of the template. The full set of elongation factors were present in these examples.

**Figure S4.**
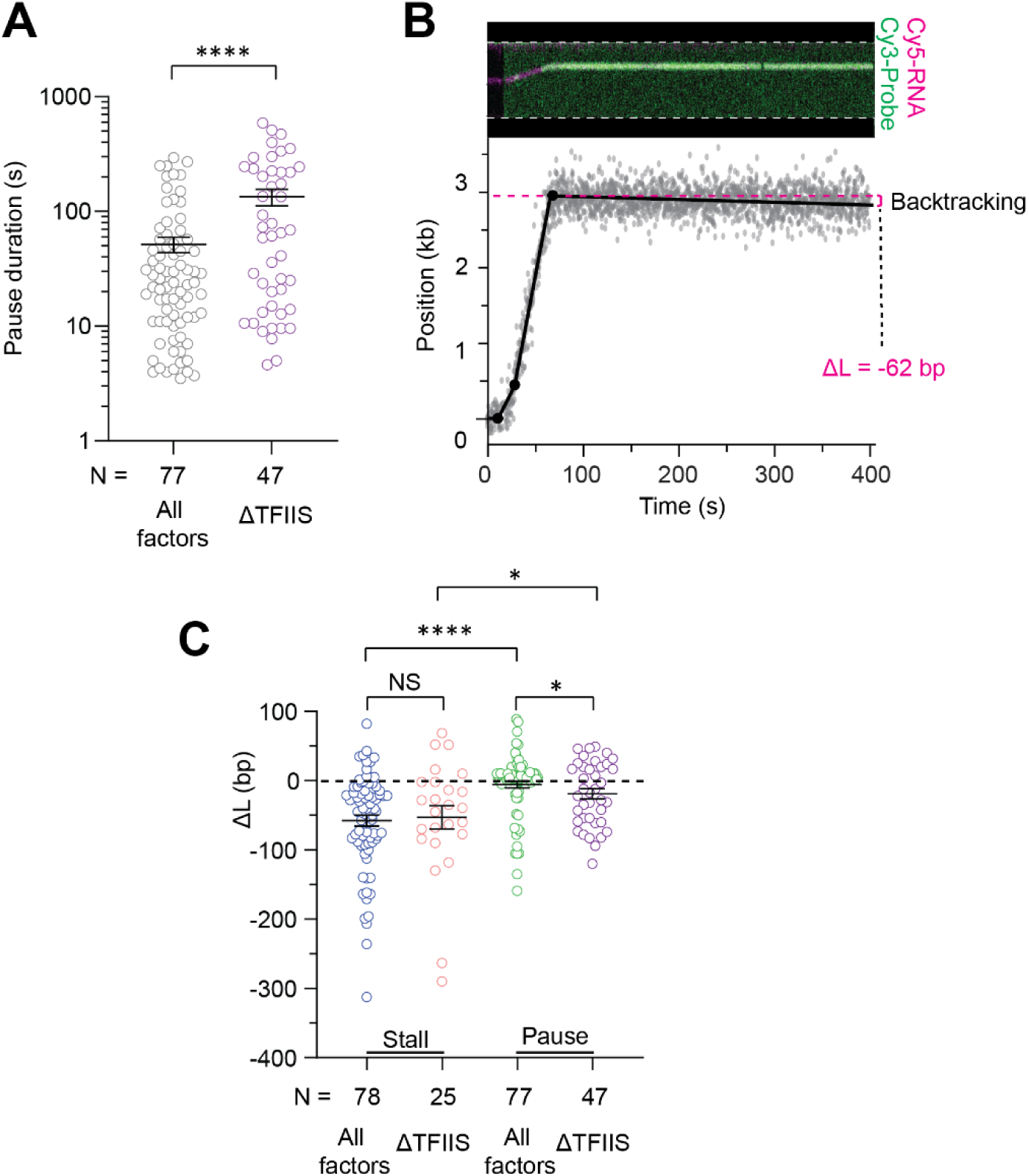
Analysis of pausing and stalling of the mammalian EC. (A) Dot plot showing the duration of pauses detected in the elongation trajectories when TFIIS was included (All factors) or omitted (ΔTFIIS). (B) An example kymograph (Top) and extracted transcription trajectory (Bottom) under the ΔTFIIS condition showing extensive EC backtracking during the stalling event. (C) Dot plot showing the position change of the EC (Δ*L*) during stalling and pausing events when TFIIS was included (All factors) or omitted (ΔTFIIS). Bars in (A) and (C) represent mean ± SEM. NS, not significant; \**P* < 0.05; \*\*\*\**P* < 0.0001. *P* values were from unpaired t tests with Welch’s correction.

**Figure S5.**
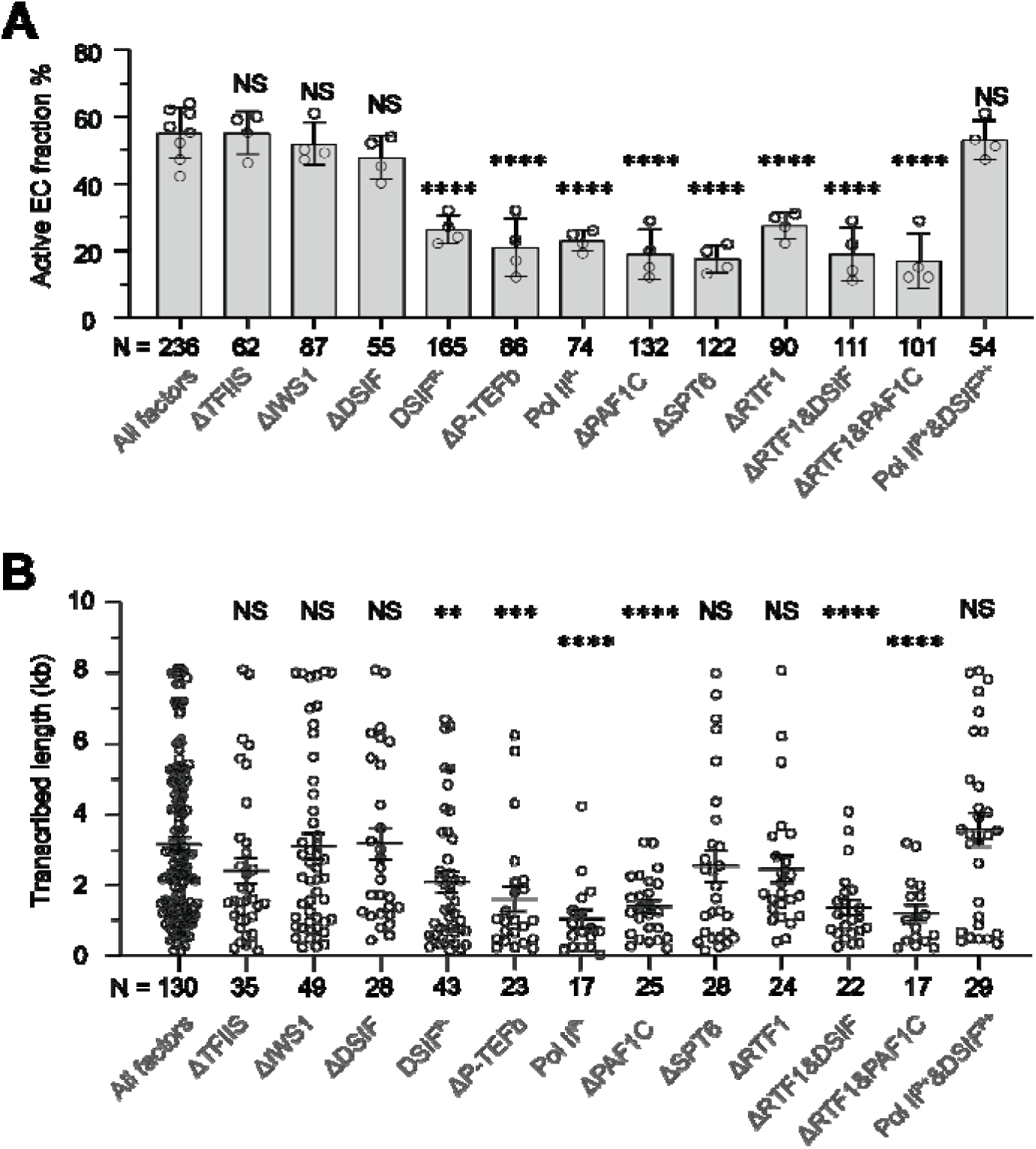
Additional data on the activity of mammalian transcription elongation under different conditions. (A) Bar plot showing the fraction of active ECs under each condition. An active EC is defined as a trajectory that exhibited detectable elongation activity (i.e. Cy3-probe signals observed.) Individual dots represent the values for each dataset from one day of data collection. *N* represent the total number of ECs (both active and inactive) for each condition collected across multiple days. (B) Dot plot showing the transcribed length before stalling for individual ECs. *N* represents the total number of active ECs analyzed for each condition. Bars represent mean ± SEM. Statistical significance was assessed by comparing each condition to the All factors condition. NS, not significant; \**P* < 0.05; \*\**P* < 0.01; \*\*\**P* < 0.001; \*\*\*\**P* < 0.0001. *P* values were from unpaired t tests with Welch’s correction.

**Figure S6.**
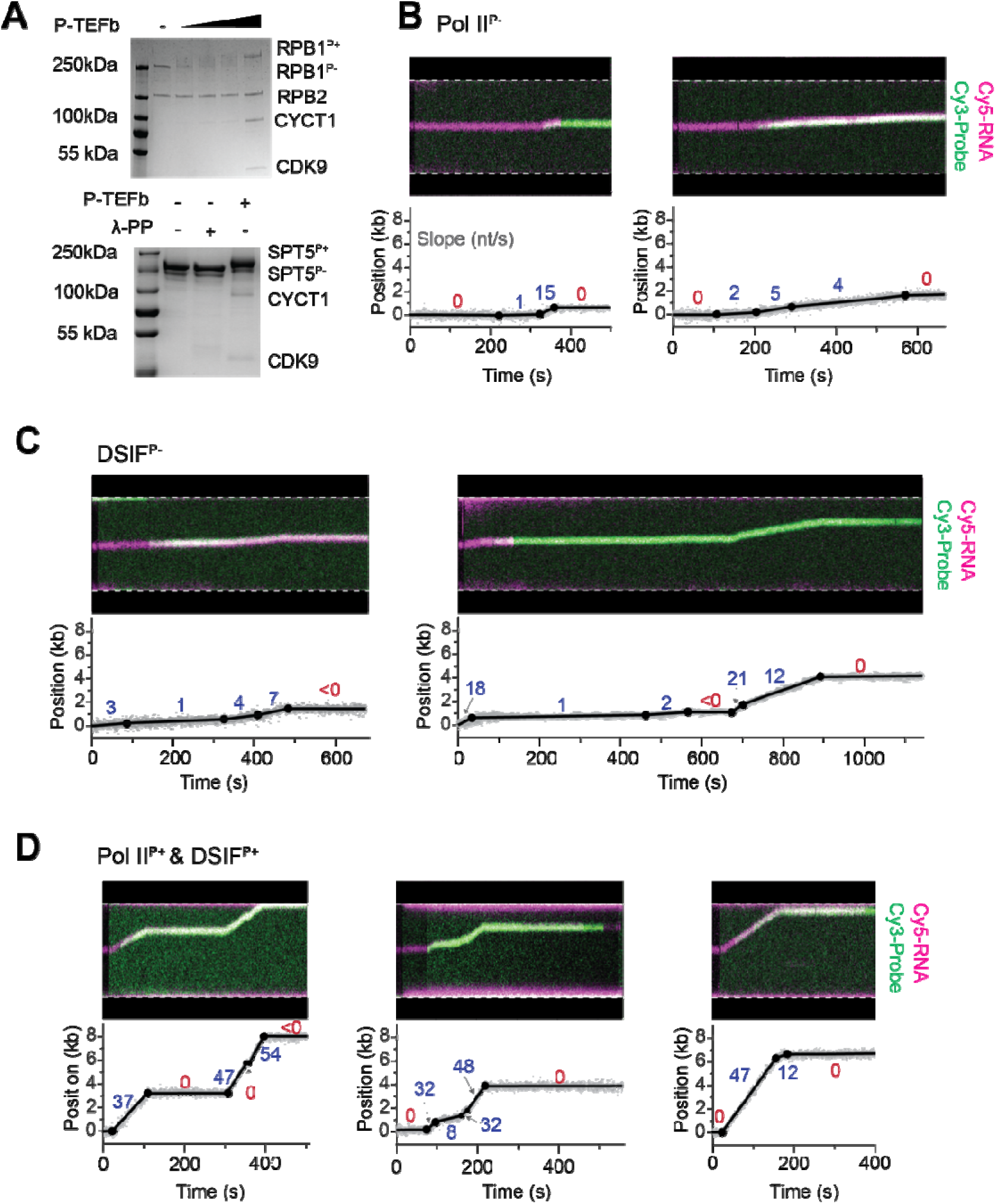
Effects of P-TEFb-mediated phosphorylation on EC activity. (A) Coomassie-stained Phos-tag SDS-PAGE gel showing phosphorylation of the RPB1 subunit of Pol II (Left) and the SPT5 subunit of DSIF (Right) by P-TEFb. Phosphorylated proteins migrated more slowly compared to their unphosphorylated version. λ-phosphatase (λ-PP) treatment confirms that SPT5 was unphosphorylated prior to the addition of P-TEFb. (B) Representative EC trajectories for the Pol II^P-^ condition. (Top) Kymographs showing the overlay of Cy5-labeled RNA (magenta) and Cy3-labeled complementary DNA probes (green) which were used to track EC progression. (Bottom) Fitted and segmented trajectories with the slope of each segment indicated. (C) Representative EC trajectories for the DSIF^P-^ condition. Details are the same as (B). (D) Representative EC trajectories for the Pol II^P+^&DSIF^P+^ condition. Details are the same as (B).

**Figure S7.**
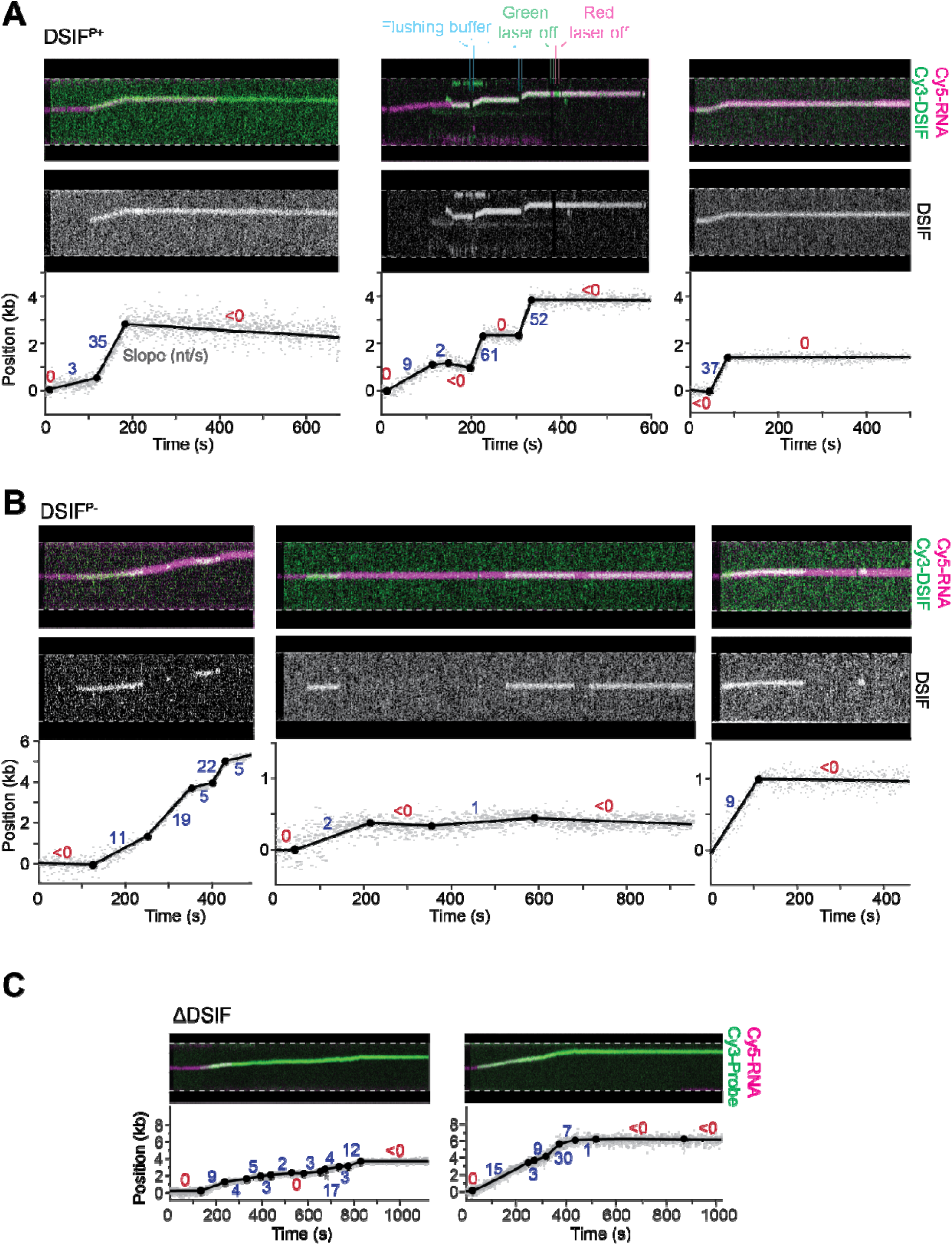
Additional data on DSIF’s role in EC activity. (A) Representative dual-color EC trajectories with Cy3-labeled DSIF and Cy5-labeled RNA. (Top) Kymographs showing the overlay of the DSIF (green) and RNA (magenta) signals. (Middle) Same kymographs but with only the DSIF signals shown. (Bottom) Fitted and segmented trajectories with the slope for each segment indicated. DSIF was phosphorylated by P-TEFb in these examples (DSIF^P+^). (B) Representative EC trajectories with unphosphorylated DSIF (DSIF^P-^). Details are the same as (A). (C) Representative kymographs (Top) and extracted trajectories (Bottom) for ECs under the condition where DSIF was omitted (ΔDSIF). Cy5-labeled RNA (magenta) and Cy3-labeled complementary DNA probes (green) were used to track EC progression.

**Figure S8.**
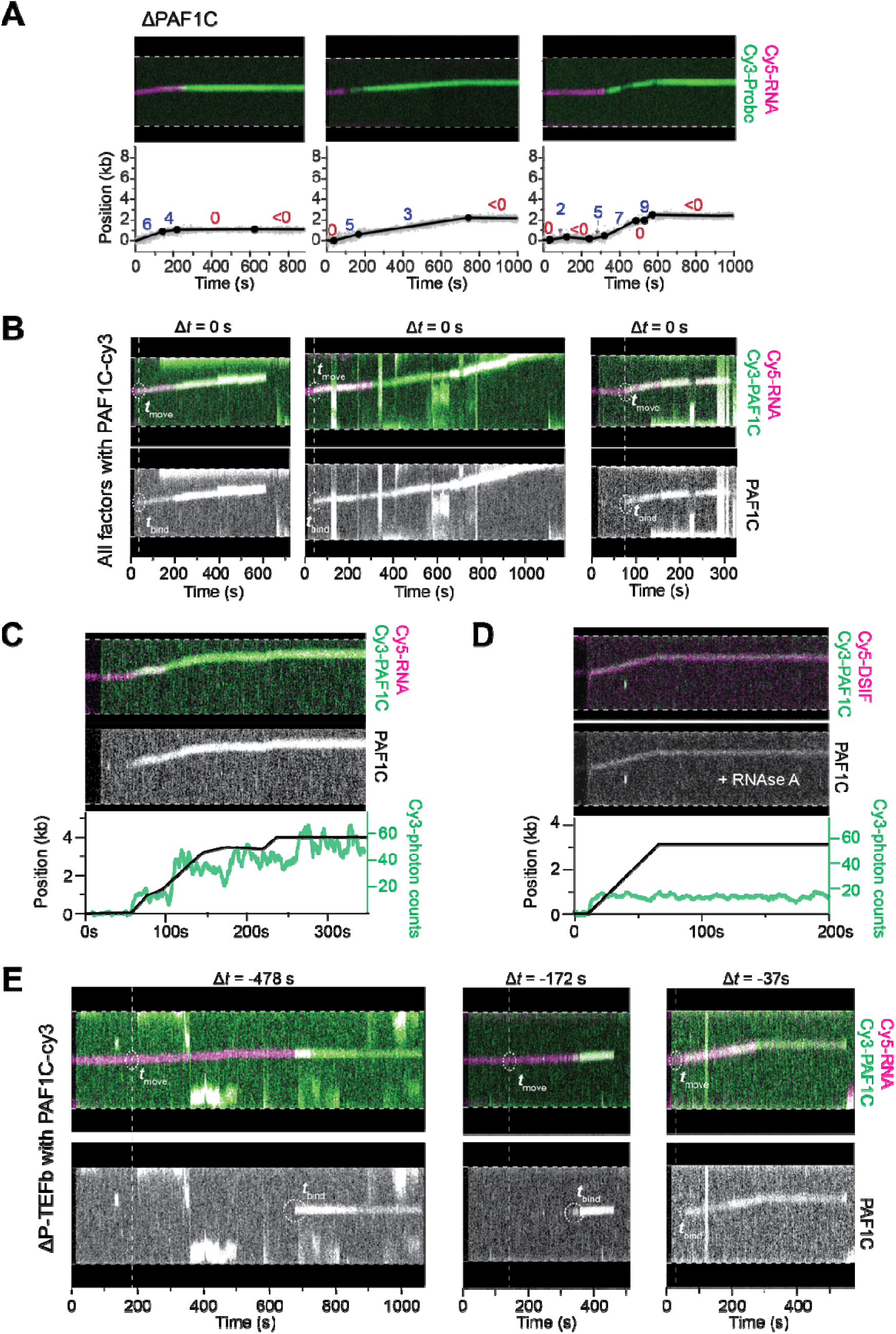
Additional data on PAF1C’s role in EC activity. (A) Representative kymographs (Top) and extracted trajectories (Bottom) for ECs under the condition where PAF1C was omitted (ΔPAF1C). Cy5-labeled RNA (magenta) and Cy3-labeled complementary DNA probes (green) were used to track EC progression. In the extracted trajectories, filled circles denote changepoints at which the trajectory was segmented. The slope for each segment is indicated. (B) Representative dual-color EC trajectories with Cy3-labeled PAF1C and Cy5-labeled RNA in the presence of all factors. (Top) Kymographs showing the overlay of PAF1C (green) and RNA (magenta) signals. *t*_move_ indicates the time point when the EC started translocation. (Bottom) The same kymographs but only showing the PAF1C signals. *t*_bind_ indicates the time point when the initial PAF1C binding to EC occurred. Δ*t* values are reported on top of the kymographs (Δ*t* = *t*_move_ – *t*_bind_). (C) An example EC trajectory showing multiple PAF1C being recruited during elongation. (Top) Kymograph showing the overlay of PAF1C (green) and RNA (magenta) signals. (Middle) The same kymograph with only the PAF1C signals shown. (Bottom) Extracted EC trajectory (black) and the corresponding Cy3 photon count (green) over time, indicating the number of PAF1C bound to EC. (D) An example EC trajectory in the presence of RNase A showing one single copy of PAF1C being recruited to the EC. (Top) Kymograph showing the overlay of Cy3-PAF1C (green) and Cy5-DSIF (magenta; in lieu of Cy5-RNA as it was digested by RNase A) signals. (Middle) The same kymograph with only the PAF1C signals shown. (Bottom) Extracted EC trajectory (black) and the corresponding Cy3 photon count (green) over time. (E) Representative dual-color EC trajectories with Cy3-labeled PAF1C and Cy5-labeled RNA under the ΔP-TEFb condition. Details are the same as (B).

**Figure S9.**
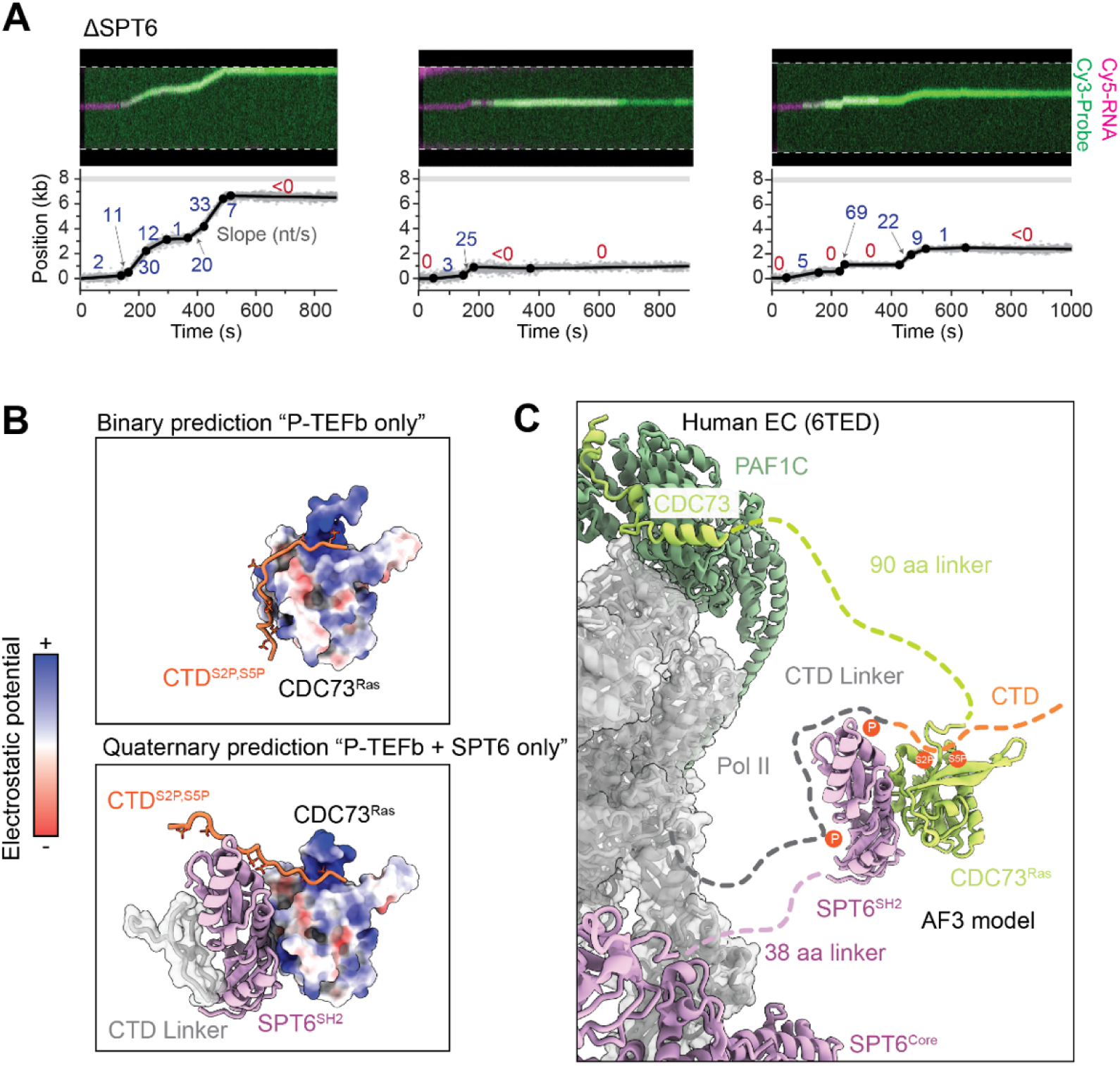
Additional data on SPT6’s role in EC activity. (A) Representative kymographs (Top) and extracted trajectories (Bottom) for ECs under the condition where SPT6 was omitted (ΔSPT6). Cy5-labeled RNA (magenta) and Cy3-labeled complementary DNA probes (green) were used to track EC progression. The slope for each segment is indicated. (B) (Top) AlphaFold 3 model of the human CDC73 Ras-like domain (CDC73^Ras^, residues: 351-531) bound to a Pol II CTD-mimicking peptide (YpSPTpSPS)_3_ (termed here CTD^S2P,S5P^). The model predicts that CTD^S2P,S5P^ (orange) interacts with CDC73^Ras^ through multiple basic patches. (Bottom) AF3 model of CDC73^Ras^ interacting with the human SPT6 SH2 domain (SPT6^SH2^, residues: 1323-1520) and CTD^S2P,S5P^. SPT6^SH2^ also interacts with the CTD linker of human RPB1 (Residues: 1523-1553). CTD^S2P,S5P^ adopts a different trajectory in this quaternary prediction compared to the binary prediction above, with one of the CDC73 basic patches now occupied by SPT6^SH2^. (C) Model for PAF1C-SPT6-Pol II interaction. Pol II, PAF1C and SPT6 chains are shown in gray, green and purple, respectively. The EC (PDB: 6TED) is shown with SPT6^SH2^-CDC73^Ras^ from the quaternary AF3 model in (B) displayed with flexible linkers connecting it to the EC. The trajectory of the CTD linker (shown as a gray dotted line with orange phosphate groups), which connects the CTD to the Pol II core, is based on how it binds SPT6^SH2^ according to a previous structural study (ref. ^10^) and in concordance with our AF3 model. The trajectory of the phosphorylated CTD (shown as an orange dotted line with orange phosphate groups) is based on the quaternary AF3 prediction in (B).

**Figure S10.**
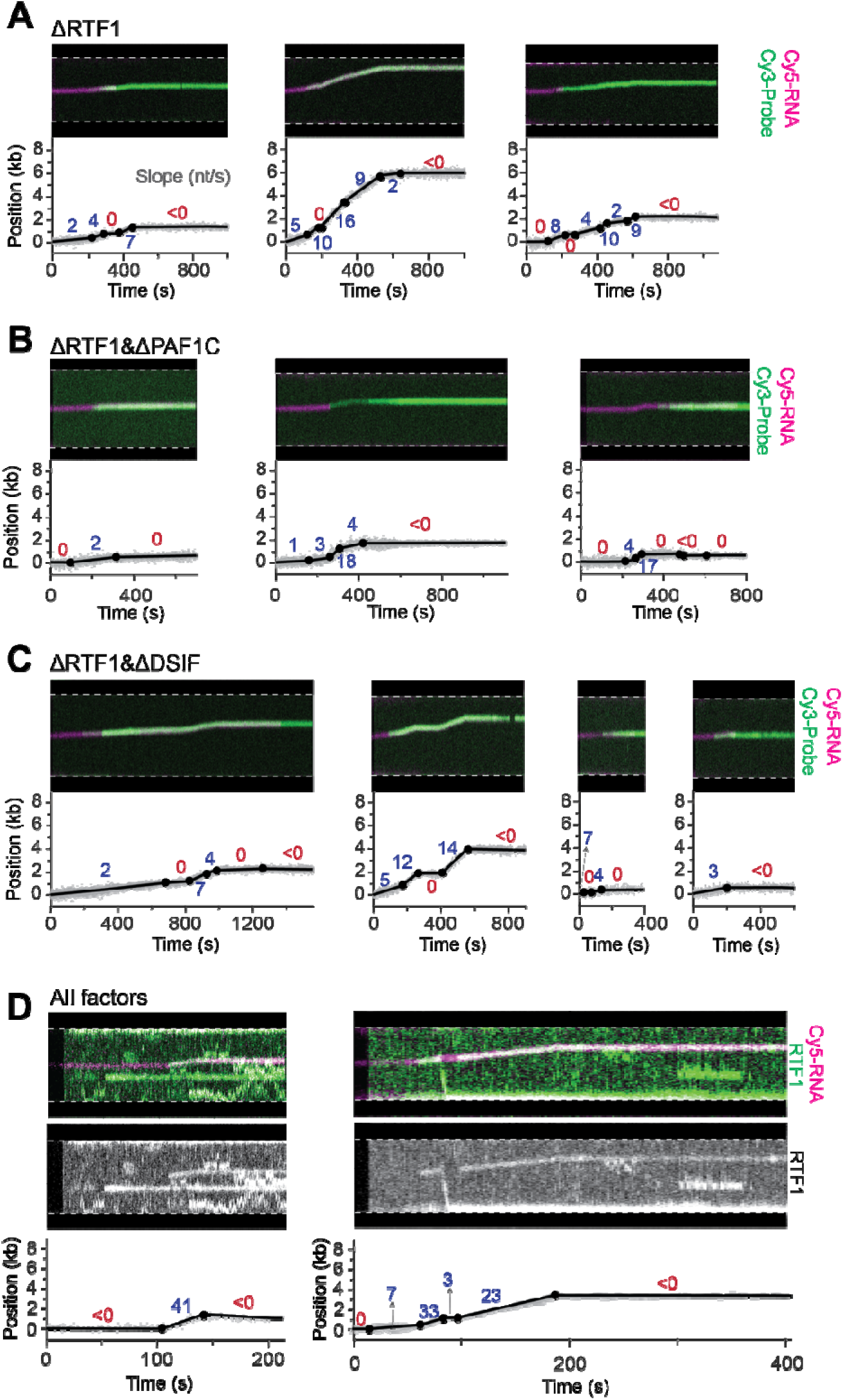
Additional data on RTF1’s role in EC activity. (A) Representative kymographs (Top) and extracted trajectories (Bottom) for ECs under the condition where RTF1 was omitted (ΔRTF1). Cy5-labeled RNA (magenta) and Cy3-labeled complementary DNA probes (green) were used to track EC progression. Filled circles in the extracted trajectories denote changepoints, dividing the trajectory into discrete linear segments. The slopes for each segment are indicated. (B) Representative kymographs (Top) and extracted trajectories (Bottom) for ECs under the condition where both RTF1 and PAF1C were omitted (ΔRTF1&ΔPAF1C). Details are the same as (A). (C) Representative kymographs (Top) and extracted trajectories (Bottom) for ECs under the condition where both RTF1 and DSIF were omitted (ΔRTF1&ΔDSIF). Details are the same as (A). (D) Representative dual-color EC trajectories with Cy3-labeled RTF1 and Cy5-labeled RNA in the presence of all factors. (Top) Kymographs showing the overlay of RTF1 (green) and RNA (magenta) signals. (Middle) Same kymographs but only showing the RTF1 signals. (Bottom) Fitted and segmented trajectories with the slope of each segment indicated.

